# Paraspeckles are constructed as block copolymer micelles through microphase separation

**DOI:** 10.1101/2020.11.02.366021

**Authors:** Tomohiro Yamazaki, Tetsuya Yamamoto, Hyura Yoshino, Sylvie Souquere, Shinichi Nakagawa, Gerard Pierron, Tetsuro Hirose

**Author notes:** Correspondence: Tomohiro Yamazaki, 1-3 Yamadaoka, Suita, Osaka 565-0871, Japan, Tetsuro Hirose, 1-3 Yamadaoka, Suita, Osaka 565-0871, Japan.

## Abstract

Paraspeckles are constructed by NEAT1_2 architectural long noncoding RNAs and possess characteristic cylindrical shapes with highly ordered internal organization, distinct from typical liquid–liquid phase-separated condensates. We experimentally and theoretically investigated how the shape and organization of paraspeckles are determined. We identified the NEAT1_2 RNA domains responsible for shell localization of the NEAT1_2 ends, which determine the characteristic internal organization. We then applied a theoretical framework using soft matter physics to understand the principles that determine the NEAT1_2 organization, shape, number, and size of paraspeckles. By treating paraspeckles as amphipathic block copolymer micelles, we could explain and predict the experimentally observed behaviors of paraspeckles upon NEAT1_2 domain deletions or transcriptional modulation. Thus, we propose that paraspeckles are block copolymer micelles assembled through microphase separation. This work provides an experimentally-based theoretical framework for the concept that ribonucleoprotein complexes (RNPs) can act as block copolymers to form RNA-scaffolding microphase-separated condensates in cells.

## Introduction

Membraneless organelles, also known as cellular bodies or biomolecular condensates, have attracted much attention because of their involvement in biological processes and pathological conditions, and because of the physical process of their formation by phase separation (Alberti and Dormann, 2019; Banani et al., 2017; Hyman et al., 2014; Shin and Brangwynne, 2017). In particular, liquid–liquid phase separation (LLPS), a type of phase separation, is widely used in a variety of biological processes (Alberti et al., 2019; Banani et al., 2017; Sabari et al., 2020; Shin and Brangwynne, 2017; Strom and Brangwynne, 2019). Many biomolecular condensates contain proteins, such as intrinsically disordered proteins and oligomer-forming proteins, and RNAs (Choi et al., 2020; Sabari et al., 2020; Sawyer et al., 2019). Although the mechanisms of protein phase separation have been extensively studied, the role of RNA remains poorly understood.

A class of RNA, termed architectural RNA (arcRNA), plays an essential scaffolding role in the formation of phase-separated condensates in various eukaryotic species from yeast to human (Chujo et al., 2016; Chujo et al., 2017; Roden and Gladfelter, 2020; Yamazaki et al., 2019; Yamazaki et al., 2018). The arcRNAs include several categories of RNA transcripts, mainly long noncoding RNAs (lncRNAs), which are pervasively transcribed from eukaryotic genomes and are important cellular regulatory factors (Quinn and Chang, 2016; Schmitt and Chang, 2017). Among the lncRNAs, non-repetitive lncRNAs, repeat-derived lncRNAs, short tandem repeat-enriched RNAs, and disease-associated repetitive RNAs have been identified as arcRNAs (Chujo et al., 2016; Chujo et al., 2017; Dumbović et al., 2018; Fox et al., 2018; Hall et al., 2017; Ninomiya and Hirose, 2020; Sasaki et al., 2009; Swinnen et al., 2019; Yap et al., 2018). NEAT1_2 lncRNA is a representative arcRNA and constructs paraspeckle nuclear bodies (Chen and Carmichael, 2009; Clemson et al., 2009; Sasaki et al., 2009; Sunwoo et al., 2009). NEAT1_2 lncRNA (22.7 kb) is a longer isoform of the *NEAT1* gene and is essential for paraspeckle formation, whereas NEAT1_1 (3.7 kb), a shorter isoform, is not essential for paraspeckle formation (Naganuma et al., 2012). NEAT1_2 lncRNAs are upregulated by several factors, including proteasome inhibition, p53 activation, and viral infections, and play critical roles in various physiological and pathological conditions (Hirose et al., 2014; Imamura et al., 2014; Mello et al., 2017; Nakagawa et al., 2014; Nakagawa et al., 2018; Standaert et al., 2014). At the molecular level in HeLa cells, a single spherical paraspeckle has been shown to contain approximately 50 NEAT1_2 molecules (Chujo et al., 2017). More than 60 paraspeckle proteins (PSPs) are enriched in paraspeckles, and several of these PSPs are required for the paraspeckle formation processes (Kawaguchi et al., 2015; Naganuma et al., 2012; Yamazaki and Hirose, 2015). Proteins, including SFPQ and NONO, are required for the expression of NEAT1_2 lncRNA, and several proteins, including NONO, FUS, and RBM14, are required for paraspeckle assembly (Hennig et al., 2015; Naganuma et al., 2012; Yamazaki et al., 2018). Specifically, oligomerization of NONO through NOPS and coiled-coil domains and interactions through the low-complexity domains of FUS and RBM14 are required for paraspeckle assembly (Hennig et al., 2015; Yamazaki et al., 2018). We have used CRISPR/Cas9-mediated dissection of NEAT1_2 in human haploid HAP1 cells to demonstrate that NEAT1_2 has modular functional RNA domains for RNA stabilization, isoform switching from NEAT1_1 to NEAT1_2, and paraspeckle assembly (Yamazaki et al., 2018). The middle domain of NEAT1_2 contains multiple binding sites for NONO and SFPQ that are necessary and sufficient for paraspeckle assembly through phase separation (Yamazaki et al., 2018). Previous studies using electron microscopy (EM) and super-resolution microscopy (SRM) have shown that paraspeckles can have a spherical or cylindrical shape (Souquere et al., 2010; West et al., 2016). The short axis (Sx) of paraspeckles is constrained (~360 nm), while the long axis (Lx) elongates upon transcriptional NEAT1_2 upregulation in cylindrical paraspeckles (Souquere et al., 2010; Yamazaki et al., 2018). Additionally, NEAT1_2 is looped and highly spatially organized within paraspeckles with the 5′ and 3′ ends of NEAT1_2 localized in the shell of the paraspeckle and the middle domain localized in the core of the paraspeckle (Souquere et al., 2010; West et al., 2016; Yamazaki et al., 2018). The 5′ and 3′ domains appear to be bundled and form distinct domains in the shells of the paraspeckles (West et al., 2016). PSPs also show specific patterns of localization within paraspeckles, suggesting a core-shell structure for the paraspeckles (West et al., 2016). These characteristic shapes and internal organization of paraspeckles are distinct from those of condensates formed by LLPS, which are usually spherical and have non-ordered internal structures. However, it is unknown how the NEAT1_2 lncRNA determines both the paraspeckle shape and the highly ordered NEAT1_2 organization within it.

Here we addressed these questions both experimentally and theoretically. We first experimentally identified the 5′ and 3′ RNA domains of NEAT1_2 that determine the shell localization by dissecting NEAT1_2 lncRNA *in vivo*. We then applied soft matter physics theories to understand the principles of the formation of the paraspeckle structure. We treated the paraspeckles as amphipathic block copolymer micelles, which form spherical and cylindrical shapes with ordered internal structures analogous to paraspeckles. Our theoretical model could explain and predict the observed behaviors of the paraspeckles formed by wild type (WT) and mutant NEAT1_2 in terms of the internal organization, size, number, and shape. Thus, this study provides a conceptual framework for the formation and structure of nuclear biomolecular condensates with RNA scaffolds.

## Results

### The 3′ terminal domain of NEAT1_2 is required for localization of the 3′ end in the shell of paraspeckles

In HAP1 WT cells, quantitative analyses using SRM have shown that the 5′ and 3′ ends of NEAT1_2 are localized in the shell of paraspeckles, as previously indicated by EM analyses (Souquere et al., 2010) (Figures 1A, F and S1A and B). It has been suggested that the 5′ and 3′ terminal domains of NEAT1_2 determine the shell localization of the 5′ and 3’ ends of the NEAT1_2 transcripts. We found that the truncated 3′ end of the NEAT1_2 Δ16.6–22.6 kb mutant (Δ3′ mutant) was localized in the core of paraspeckles, even in the presence of the 3′-terminal triple-helix structure (Yamazaki et al., 2018). We have also observed that the expression of NEAT1_1 and NEAT1_2 in Δ3′ mutant cells was comparable to that in WT cells (Yamazaki et al., 2018). Then, we further characterized this Δ3′ mutant. In Δ3′ mutant cells, in which no paraspeckle assembly defects were detected (Figure S1C), quantitative analysis using SRM showed the localization of the 3′ end of the Δ3′ mutant to the core of the paraspeckles, whereas the 5′-end remained localized in the shell (Figures 1A, B, and F and S1B). The EM analysis confirmed that the truncated 3′ end of the Δ3′ mutant was localized in the core without affecting the localization of the 5′ end in the shell (Figures 1C, F and S1D). 5′ and D2 probes for EM were used to detect the localization in the shells and cores, respectively, in WT cells, as previously reported (Souquere et al., 2010) (Figure S1E and F). Thus, the 16.6–22.6 kb region of NEAT1_2 is required for the shell localization of the 3′ end of NEAT1_2, but its deletion does not appreciably affect the assembly of the paraspeckles.

**Figure 1.**
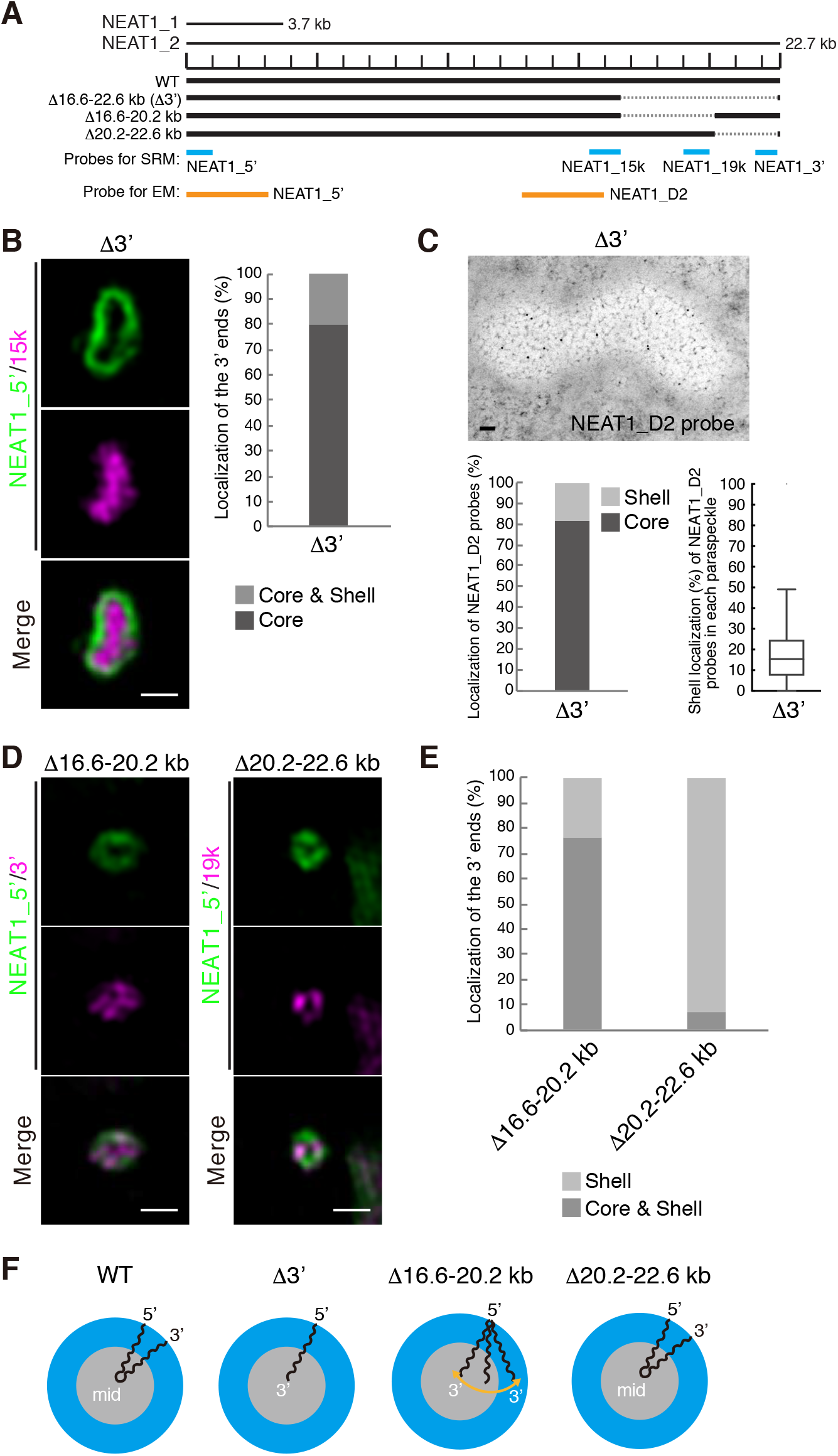
Deletion of the 3′ terminal domain of NEAT1_2 causes core localization of the 3′ ends of NEAT1_2 within the paraspeckle. (A) The schematics of human NEAT1_2 (WT) and the mutants with deletions in the 3′ terminal regions. The NEAT1 transcripts are shown above with a scale. The gray dashed lines represent the deleted regions. The positions of NEAT1 probes used in SRM (blue) and EM (orange) are shown. (B) (left) SRM images of paraspeckles in HAP1 NEAT1 Δ3′ mutant cells (Δ3′) detected by NEAT1_5′ (green) and NEAT1_15k (magenta) FISH probes in the presence of MG132 (5 μM for 6 h). Scale bar, 500 nm. (right) Graph showing the proportion of paraspeckles with localization of the NEAT1 3′ ends in the core or in the core and shell (n = 44). (C) (upper) EM observation of the paraspeckles in Δ3′ cells using NEAT1_D2 probes in the presence of MG132 (5 μM for 17 h). Scale bar, 100 nm. (lower, left) Graph showing the proportion of localization of the NEAT1 region detected by NEAT1_D2 probe (248 gold particles) within the paraspeckles in Δ3′ cells. (lower, right) Graph showing the proportion of localization of NEAT1_D2 probes in each paraspeckle in Δ3′ cells (n = 17). (D) SRM images of the paraspeckles in Δ16.6–20.2 kb and Δ20.2–22.6 kb cells detected by NEAT1_5′ (green) and NEAT1_3′ or 19k (magenta) FISH probes in the presence of MG132 (5 μM for 6 h). Scale bar, 500 nm. (E) Graph showing the proportion of paraspeckles with localization of the NEAT1 3′ ends to the core and shell or the core in Δ16.6–20.2 kb and Δ20.2–22.6 kb cells treated with MG132 (5 μM for 6 h). (Δ16.6–20.2 kb: n = 230, Δ20.2–22.6 kb: n = 99) (F) Schematics of the NEAT1_2 configuration in WT and deletion mutants.

To determine the precise region of the NEAT1_2 domain that is required for the shell localization, we established two HAP1 mutant cell lines (Δ16.6–20.2 kb and Δ20.2–22.6 kb) (Figure 1A). In both cell lines, the NEAT1_2 expression levels were comparable to that of the WT, and the paraspeckles were formed similarly to the WT (Figure S1C and G). As shown by SRM, in the majority of the Δ16.6–20.2 kb cells, the 3′ ends of this NEAT1_2 mutant were distributed in both the core and shell of the paraspeckles (Figure 1D and E). In contrast, the 3′ ends of NEAT1_2 in the Δ20.2–22.6 kb cells were mostly detected in the shell, similar to the WT paraspeckles, indicating that this region played only a minor role in the shell localization (Figure 1D and E). Altogether, these data revealed that, unlike the WT and Δ3′ mutant, the Δ16.6–20.2 kb mutant showed a random distribution of the 3′ end within the paraspeckles (Figure 1F); thus, the NEAT1_2 16.6–20.2 kb region has a major role in the proper shell localization of the 3′ end of NEAT1_2.

### The 5′ terminal domain of NEAT1_2 is required for shell localization of the 5′ end within the paraspeckles

We next investigated whether the 5′ terminal domain of NEAT1_2 has a role in the shell localization of the 5′ end. We used two mutant cell lines, the previously established NEAT1 Δ0–0.8 kb and a newly established Δ0–1.9 kb (Δ5′ mutant), in which NEAT1_2 was expressed comparably to that in WT cells, although a NEAT1 0–1 kb deletion has been previously shown to prevent NEAT1_2 accumulation (Yamazaki et al., 2018) (Figures 2A and S2A). No paraspeckle assembly defects were observed in these cell lines (Figure S2B). Our SRM analysis showed that the signals of the 5′ end were randomly localized within the paraspeckles for both mutants, whereas the signals of the 3′ end were detected in the shells in both cell lines (Figures 2B and C and S2C). Consistent with these data, EM observations confirmed the random localization of the 5′ region of NEAT1_2 in the Δ5′ mutant cells (Figure 2D). We also examined the localization of the middle region of NEAT1_2 in detail using the D2 probe for EM (Figure 2A). The middle region was mainly detected in the inner and middle layers of the paraspeckles in WT cells, as has been previously reported (Souquere et al., 2010), but in the Δ5′ mutant cells, the middle region was mainly detected in the outer and middle layers of the paraspeckles, suggesting that this region had a tendency toward being localized in the shells of the paraspeckles (Figure S2D). Collectively, these data showed that the 5′ terminal domain of NEAT1 is required for the shell localization of the 5′ end and influences the internal distribution of the NEAT1_2 transcripts within the paraspeckles (Figure 2E).

**Figure 2.**
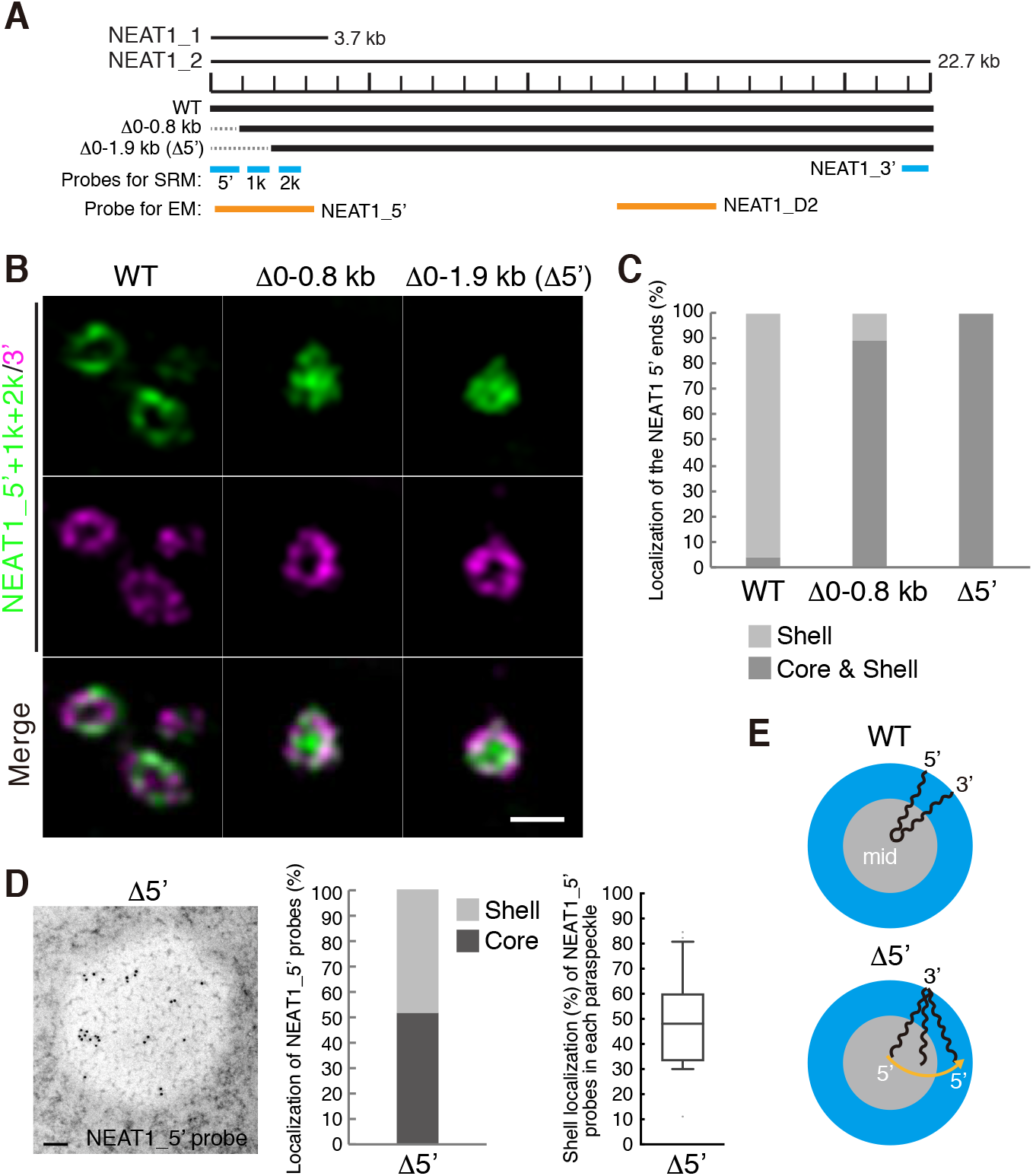
Deletion of 5′ terminal domain of NEAT1_2 causes random distribution of the NEAT1_2 5′ ends within the paraspeckle. (A) The schematics of WT NEAT1_2 and mutants with deletions in the 5′ terminal regions are shown in Figure 2A. The positions of NEAT1 probes used in SRM (blue) and EM (orange) are shown. (B) SRM images of the paraspeckles in Δ0–0.8kb and Δ0–1.9kb (Δ5′) cells treated with MG132 (5 μM for 6 h) detected by NEAT1_5′, 1k, and 2k (green) and NEAT1_3′ (magenta) FISH probes. Scale bar, 500 nm. (C) Graph showing the proportion of paraspeckles with localization of the NEAT1 5′ ends to the core and shell or the shell in WT, Δ0–0.8 kb, and Δ5′ cells treated with MG132 (5 μM for 6 h). (WT: n = 159, Δ0–0.8 kb: n = 54, Δ5′: n = 48) (D) (left) EM observation of the paraspeckles in Δ5′ cells treated with MG132 (5 μM for 17 h) using NEAT1_5′ probes. Scale bar, 100 nm. (middle) Graph showing the proportion of localization of NEAT1_5′ probes (347 gold particles) within the paraspeckles in Δ5′ cells. (right) Graph showing the proportion of the localization of NEAT1_5′ probes in each paraspeckle in Δ5′ cells (n = 20). (E) Schematics of the NEAT1_2 configuration in the WT and Δ5′ mutant.

As the truncated 3′ terminal region of NEAT1_2 was localized in the paraspeckle core in the Δ3′ mutant cells, we attempted to obtain the corresponding 5′ terminal deletion clone of NEAT1 with exclusive core localization of the 5′ end, by establishing the additional 5′ terminal deletion cell line, NEAT1 Δ0–2.8 kb. However, in this cell line, NEAT1_2 was not expressed at all; hence, paraspeckles were not formed (Figure S2E and F), precluding further analysis.

NEAT1_1 overlaps with the 5′ terminal region of NEAT1_2 and it is impossible to distinguish these two transcripts by fluorescence *in situ* hybridization (FISH). Thus, to validate the configuration of the 5′ end of NEAT1_2 in the Δ5′ mutant cells, we established the Δ5′/ΔPAS cell line, which lacks a polyadenylation signal (PAS) for NEAT1_1 production. In this cell line, NEAT1_1 expression was reduced (Figure S2A). The localization of the 5′ region of NEAT1_2 was still random within the paraspeckles, as observed in the Δ5′ mutant cells (Figure S2G), suggesting that NEAT1_1 did not affect the arrangement of NEAT1_2 observed above. Thus, 5′ terminal deletion triggers the internalization of the 5′ truncated end of NEAT1_2.

### Simultaneous deletion of the 5′ and 3′ terminal domains of NEAT1_2 causes random distribution of NEAT1_2 within the paraspeckles

As we had identified the NEAT1_2 RNA domains required for the shell localization of the 5′ and 3′ regions of NEAT1_2, we next investigated how the deletion of both the domains, 0–1.9 kb and 16.6–22.6 kb, influenced the NEAT1_2 spatial organization. We established a Δ5′/Δ3′ (Δ0–1.9 kb/Δ16.6–22.6 kb) mutant cell line, in which NEAT1_2 was expressed comparable to that in WT cells and no paraspeckle assembly defects were observed (Figures 3A and S3A and B). In contrast to the highly ordered NEAT1_2 organization of the paraspeckles in WT cells, SRM observations clearly showed a random distribution of the 5′ and 3′ regions of NEAT1_2 in the Δ5′/Δ3′ mutant cells (Figure 3B and C). Additionally, EM analyses revealed that the 5′ terminal region was detected equally in both the shell and core of the paraspeckles, while the 3′ terminal region was almost randomly distributed with a slight tendency toward core localization (Figure 3D). These data were confirmed by SRM of the Δ5′/Δ3′/ΔPAS mutant cell line, which has reduced NEAT1_1 expression compared with the WT (Figure S3C and D). Together with the data shown in Figures 1 and 2, these data suggested that both the 5′ and 3′ domains were essential for localizing both the ends of NEAT1_2 in the shell of paraspeckles.

**Figure 3.**
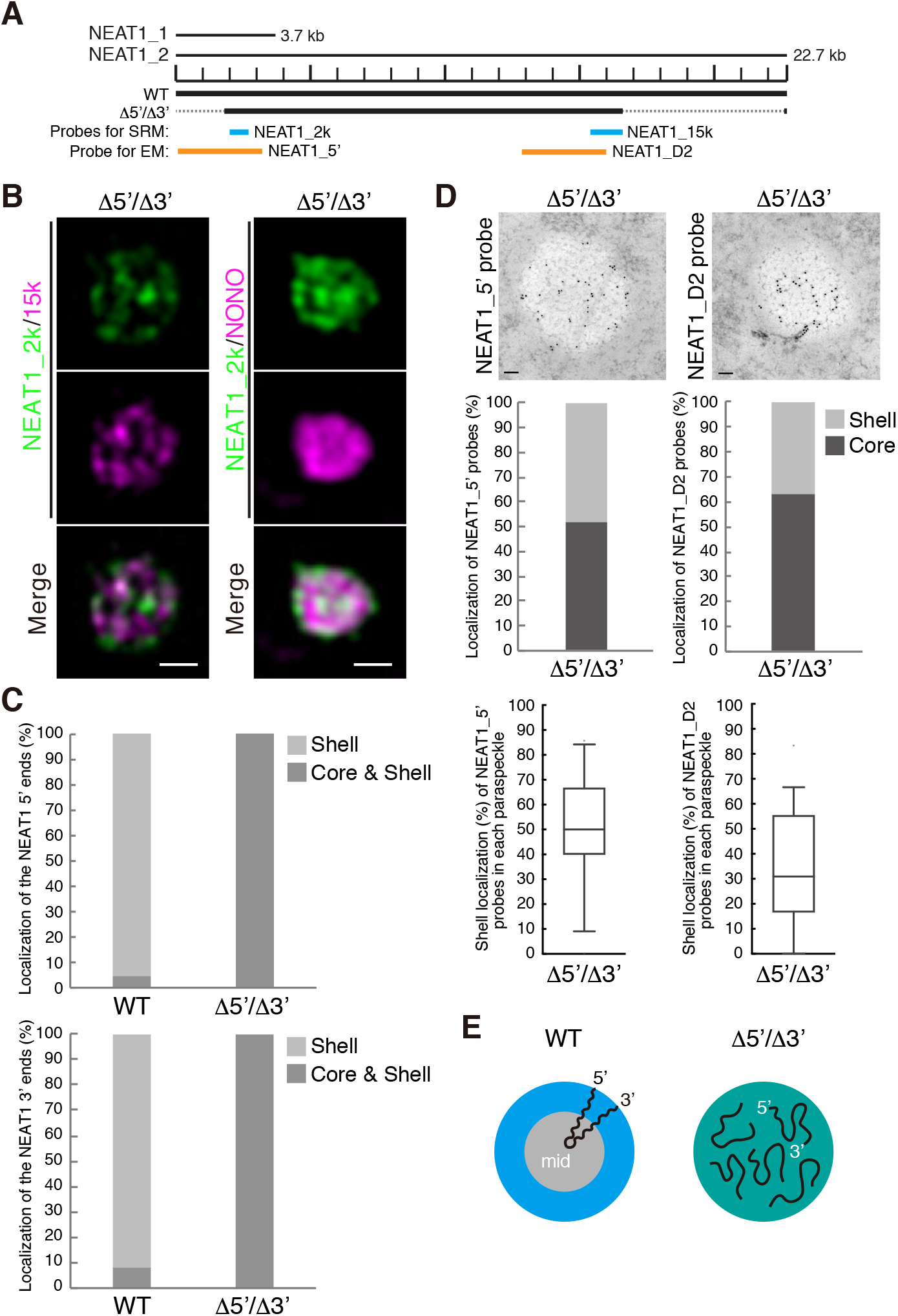
Deletion of both the 5′ and 3′ terminal domains of NEAT1_2 causes random distribution of both the 5′ and 3′ ends of NEAT1_2 within the paraspeckle. (A) The schematics of the WT and the mutant (Δ5′/Δ3′) with deletion of the 5′ and 3′ terminal regions are shown as Figure 1A. The positions of the NEAT1 probes used in SRM (blue) and EM (orange) are shown. (B) SRM images of the paraspeckles in MG132-treated (5 μM for 6 h) HAP1 NEAT1 Δ5′/Δ3′ cells detected by NEAT1_2k (green) and NEAT1_15k FISH probes (left, magenta) or NONO immunofluorescence (IF) (right, magenta). Scale bar, 500 nm. (C) Graph showing the proportion of paraspeckles with localization of the NEAT1 5′ (upper; WT: n = 41, Δ5′/Δ3′ kb: n = 44) and 3′ (lower: WT, n = 167, Δ5′/Δ3′ kb: n = 34) ends to the core and shell or the shell in WT and Δ5′/Δ3′ kb cells treated with MG132 (5 μM for 6 h). (D) (upper) EM observation of the paraspeckles in MG132-treated (5 μM for 17 h) Δ5′/Δ3′ cells using NEAT1_5′ (left) and NEAT1_D2 probes (right). Scale bar, 100 nm. (middle) Graph showing the proportion of the localization of NEAT1_5′ (left, 310 gold particles) and NEAT1_D2 (right, 294 gold particles) probes within the paraspeckles in HAP1 NEAT1 Δ5′/Δ3′ cells. (lower) Graph showing the proportion of the localization of NEAT1_5′ (left, 19 cells) and D2 (right, 22 cells) probes in each paraspeckle in Δ5′/Δ3′ cells. (E) Schematics of the NEAT1_2 configuration in the WT and Δ5′/Δ3′ mutant.

### Block copolymer micelle model of paraspeckles

We have recently established the framework for arcRNA-induced phase separation using an extension of the Flory-Huggins theory, which is the standard theory for the phase separation of polymers in solutions (Yamamoto et al., 2020a). In this framework, arcRNA-induced, phase-separated assemblies are treated as disordered condensates formed through macroscopic phase separation, such as LLPS. However, this premise does not fit the nature of paraspeckles. Paraspeckles possess highly ordered internal structures and their shapes can be cylindrical, as well as spherical, which is distinct from LLPS-induced spherical droplets (Souquere et al., 2010; West et al., 2016). The 5′ and 3′ terminal domains compose the shell, whereas, the middle domain, which is the primary paraspeckle assembly domain that interacts with oligomer-forming NONO proteins, composes the core of paraspeckles (Souquere et al., 2010; West et al., 2016; Yamazaki et al., 2018). The core-shell structure is analogous to the micelle structure of block copolymers, that is, polymers composed of two or more chemically distinct polymer blocks (Bates and Bates, 2017; Mai and Eisenberg, 2012). The simplest AB diblock copolymers, which are composed of two distinct polymer blocks, self-assemble to form various shapes, including spheres and cylinders (Figure 4A). These shapes are determined by i) the ratio of the lengths of the A and B blocks; ii) the interaction strength between the monomer units in the A and B blocks; and iii) the polymer concentration (Bates and Bates, 2017; Mai and Eisenberg, 2012). The size of the phases depends on the length of the block copolymer because the A and B blocks are connected and cannot separate from each other. Such phase separation, in which the assemblies have an optimal size and shape, is called microphase separation (Mai and Eisenberg, 2012). Amphipathic block copolymers, in which two or more blocks with different affinities to a solvent (e.g., hydrophilic and hydrophobic blocks) are connected, self-assemble into spherical and cylindrical micelles in water (Bates and Bates, 2017; Mai and Eisenberg, 2012). The characteristics of paraspeckles, such as the cylindrical shape and the ordered internal organization of NEAT1_2, are reminiscent of block copolymer micelles. Because NEAT1_2 forms RNPs with various RNA-binding proteins (RBPs), in which each of 5′ and 3′ terminal domains and the middle domain all form distinct RNP domains within a paraspeckle (Naganuma et al., 2012; Nakagawa et al., 2018; West et al., 2016), we considered NEAT1_2 RNPs to be analogous to ABC triblock copolymers (Moughton et al., 2012) (Figure 4B). The A and C blocks corresponding to the 5′ and 3′ terminal regions, respectively, of NEAT1_2 outside of the major middle domain (8–16.6 kb) can be treated as hydrophilic domains exposed to the nucleoplasm. The B block, that is, the middle domain of NEAT1_2 that NONO proteins interact with and oligomerize to bridge NEAT1_2 to induce paraspeckle assembly with associated proteins, such as FUS (Hennig et al., 2015; Naganuma et al., 2012; Yamazaki et al., 2018), can be treated as the hydrophobic core domain of a paraspeckle (Figure 4B). We then constructed a theoretical model of paraspeckle formation by treating paraspeckles as ABC triblock copolymer micelles (Yamamoto et al., 2020b). In this model, we treated paraspeckles as spherical micelles and each block was considered to be composed of identical units for simplicity (also see the Discussion section). We used this model to analyze the localization of the A (and C) blocks, corresponding to the 5′ (and 3′) ends of NEAT1_2, and the size of the micelles (paraspeckles) (Figure 4C). A fraction, α of the A blocks are localized in the shell, and the other fraction, 1–α, are localized in the core. The fraction α is determined by minimizing the free-energy of the micelle, which takes into account: i) the surface free energy (which is the energy cost because the B block units at the surface of the core cannot bind with other B block units, unlike the units in the interior of the core, and is proportional to the surface area); ii) the excluded-volume interactions between the A and B blocks in the core (which is the energy cost because the A block units in the core disturb the binding between the B block units); iii) the excluded-volume interactions between the A blocks, and between the C blocks, in the shell (which are repulsive interactions between A or C blocks because these blocks are hydrophilic and thus tend to mix with the solvent); and iv) the mixing free energy (which is the contribution of thermal fluctuations to randomly distribute the A blocks between the core and the shell) (Doi 1996; Yamamoto et al., 2020b). Theoretical analysis of the localization of A (and C) blocks [the 5′ (and 3′) terminal domains of NEAT1_2] could rationalize the phenotypes observed in the NEAT1_2 deletion mutants described above: As the length of the A or C block becomes shorter, the A or C block is redistributed to the core of the micelles (paraspeckles) (Figure 4D). Therefore, our micelle model can be used to explain the reorganization of the NEAT1_2 ends within the paraspeckles as observed in the NEAT1_2 mutants.

**Figure 4.**
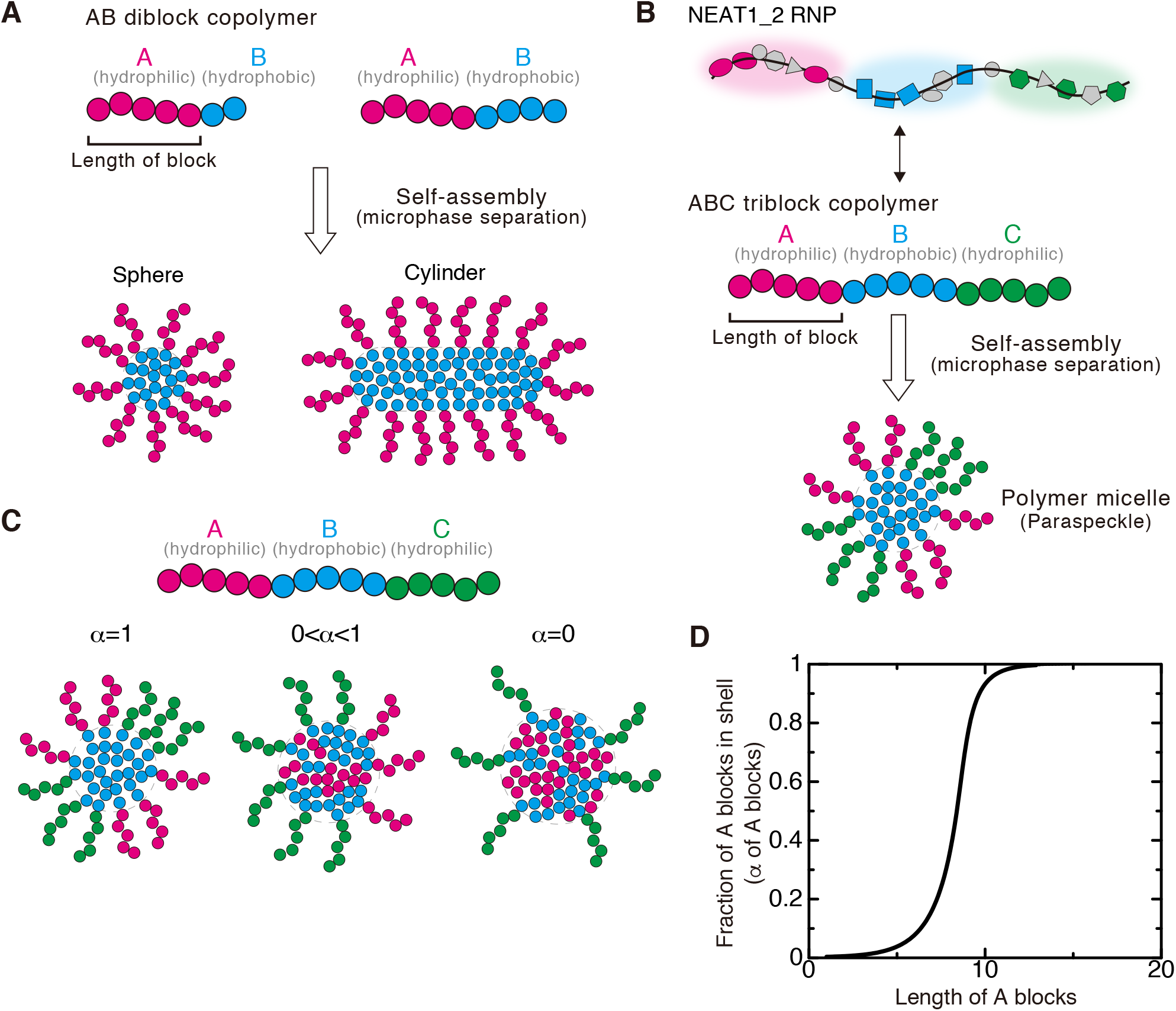
Block copolymer micelle model of paraspeckles (A) Schematics of AB diblock copolymers with different lengths of blocks and examples of the microphases (sphere and cylinder) that they form. Different blocks are shown in different colors (magenta and cyan). (B) Schematics showing analogies between NEAT1_2 RNP (upper) and ABC triblock copolymer (middle), and a triblock copolymer micelle and a paraspeckle (lower). Different blocks consisting of RNA and the putative partner RBPs are shown in different colors (magenta, cyan, and green). (C) In the copolymer micelle model, B blocks are localized in the core and C blocks are localized in the shell. A fraction, α of A blocks are localized in the shell, and the other fraction, 1–α, are localized in the core. (D) The fraction α of the A blocks in the shell versus the length of the A blocks (represented by the number of segments) of spherical paraspeckles in the steady state. The parameters used for the calculations are shown in the STAR Methods.

### Paraspeckles behave as triblock copolymer micelles

We further attempted to validate our theoretical model experimentally. The model predicts that the organization within microphase-separated condensates will influence the size of the assemblies. In this model, based on the theory of block copolymers (Halperin and Alexander, 1989; Semenov et al., 1995; Zhulina et al., 2005), the size of the paraspeckles is constrained by the repulsive excluded-volume interactions between the A blocks, or between the C blocks, localized in the shell of the paraspeckles because these interactions limit the number of the incorporated NEAT1_2 molecules and the size of the paraspeckles (Yamamoto et al., 2020b). Without these repulsive interactions, as in the case of condensates assembled by LLPS, the shape and dynamics of the condensates are governed by the surface tension; condensates are therefore always spherical and grow via coalescence to minimize the surface area (Doi, 2013). The fact that paraspeckles of WT NEAT1_2 can form cylinders and be present as clusters, instead of showing coalescence (Hirose et al., 2014; Souquere et al., 2010; Visa et al., 1993; West et al., 2016), suggested the repulsive excluded-volume interactions between the 3′ terminal regions, and those between the 5′ terminal regions, are analogous to the interactions in polymer micelles (Halperin and Alexander, 1989; Semenov et al., 1995; Zhulina et al., 2005). The deletion of the 5′ and/or 3′ terminal domains of NEAT1_2, corresponding to the A and/or C blocks, reduced the excluded-volume interactions (which decreased with a decrease in the local concentration of the A and/or C blocks) because of the redistribution of the terminal domains to the core (Figure 4D). Then, the number of incorporated NEAT1_2 molecules was increased, leading to the enlargement of the paraspeckles. Our model therefore predicts that, as the length of the A or C blocks decreases, both the size of the paraspeckles and the number of incorporated NEAT1_2 molecules become larger (Figure 5A and B).

**Figure 5.**
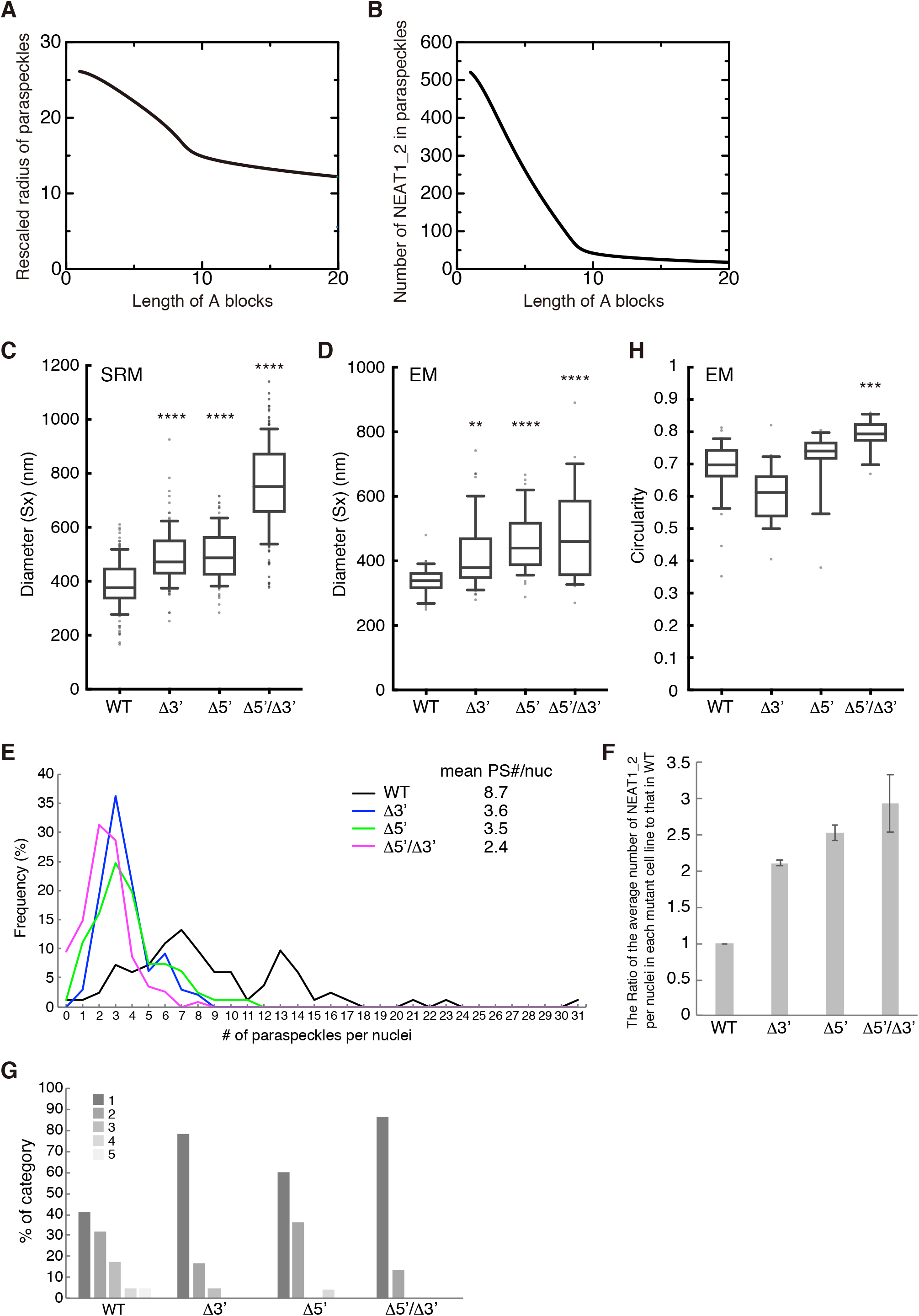
Paraspeckles behave as triblock polymer micelles. (A) Theoretical calculation of the radius of paraspeckles versus the length of the A blocks (represented by the number of segments) of spherical paraspeckles in the steady state. The radius was rescaled by the segment length. The parameters used for the calculations are shown in the STAR Methods. (B) Theoretical calculation of the number of NEAT1_2 transcripts versus the length of the A blocks (represented by the number of segments) of spherical paraspeckles in the steady state. The parameters used for the calculations are shown in the STAR Methods. (C) Diameters (Sx) of the paraspeckles in WT, Δ3′, Δ5′, and Δ5′/Δ3′ cells treated with MG132 (5 μM for 6 h) determined by SRM (WT: n = 129, Δ3′: n = 106, Δ5′: n = 100, Δ5′/Δ3′: n = 292). WT (mean size: 387.5 nm), Δ3′ (mean size: 492.4 nm), Δ5′ (mean size: 497.7 nm), and Δ5′/Δ3′ mutant (mean size: 753.3 nm). (****: *P* < 0.0001, compared with WT). (D) Diameter (Sx) of the paraspeckles in WT, Δ3′, Δ5′, and Δ5′/Δ3′ cells treated with MG132 (5 μM for 17 h) determined by EM (WT: n = 35, Δ3′: n = 49, Δ5′: n = 39, Δ5′/Δ3′: n = 28). WT (mean size: 339.4 nm), Δ3′ (mean size: 416.4 nm), Δ5′ (mean size: 457.4 nm), and Δ5′/Δ3′ (mean size: 488.9 nm). (**: *P* = 0.0041, ****: *P* < 0.0001, compared with WT). (E) The number of paraspeckles per nuclei in each cell line. The mean numbers are shown (mean PS (paraspeckle) #/nuc) (WT: n = 83, Δ3′: n = 99, Δ5′: n = 103, Δ5′/Δ3′: n = 115). Statistical analysis showed a significant reduction in the paraspeckle numbers in the Δ3′ (*P* < 0.0001), Δ5′ (*P* < 0.0001), and Δ5′/Δ3′ (*P* < 0.0001) mutants compared with the WT. The number of paraspeckles in the Δ5′/Δ3′ was significantly fewer than that in the Δ3′ and Δ5′ (*P* < 0.0001 compared with Δ3′, *P* = 0.0023 compared with Δ5′). (F) The ratio of the average number of NEAT1_2 per nuclei in each mutant cell line to that in the WT. Data are represented as mean ± SD (n = 3). (G) The proportion of the number of the paraspeckles in each cluster determined by EM in HAP1 WT, Δ3′, Δ5′, and Δ5′/Δ3′ cells treated with MG132 (5 μM for 17 h). (WT: n = 82, Δ3′: n = 37, Δ5′: n = 53, Δ5′/Δ3′: n = 34). (H) Circularity [4π x surface area/(perimeter)^2^] of the paraspeckles in HAP1 WT, Δ3′, Δ5′, and Δ5′/Δ3′ cells treated with MG132 (5 μM for 17 h) determined by EM (WT: n = 33, Δ3′: n = 21, Δ5′: n = 19, Δ5′/Δ3′: n = 19). WT (mean: 0.6844), Δ3′ (mean: 0.6057), Δ5′ (mean: 0.7151), Δ5′/Δ3′ (mean: 0.7921). (***: *P* = 0.0001, compared with WT).

To experimentally examine this prediction, we measured the sizes of the paraspeckles in the WT, Δ3′, Δ5′, and Δ5′/Δ3′ mutant cell lines. Measurements using SRM showed that the Sx values of the paraspeckles in the Δ3′ and Δ5′ mutants were larger than that in the WT (Figure 5C). The Sx value of the paraspeckles in the Δ5′/Δ3′ mutant was much larger than those in the WT and the other mutants, although the length of the NEAT1_2 Δ5′/Δ3′ arcRNA itself was the shortest (Figure 5C). We validated these data using EM and obtained similar results (Figure 5D). The Sx values of all the deletion mutants, Δ3′, Δ5′, and Δ5′/Δ3′, were larger than that of the WT, although the Sx value of the Δ5′/Δ3′ mutant may have been underestimated because of the spherical shape of the paraspeckles (as described below). Additionally, the Lx values of the paraspeckles in the Δ3′, Δ5′, and Δ5′/Δ3′ cells tended to be larger compared with those of the paraspeckles in the WT cells (Figure S4A). The surface areas of the paraspeckles, as observed by EM, were larger in the Δ3′, Δ5′, and Δ5′/Δ3′ cells than those of the paraspeckles in the WT cells (Figure S4B). We also experimentally examined whether more NEAT1_2 RNA molecules were incorporated into the paraspeckles in the Δ3′, Δ5′, and Δ5′/Δ3′ cells than into those in the WT cells as theoretically predicted. The NEAT1_2 expression levels were quantified by RT-qPCR and the paraspeckle number per nuclei was determined by high-resolution microscopy in these cell lines. Because NEAT1_2 arcRNAs are exclusively localized in paraspeckles (Chujo et al., 2017; Sasaki et al., 2009), we estimated the number of NEAT1_2 molecules per paraspeckle, relative to the value in the WT, as the expression level divided by the average number of paraspeckles per nuclei. The analysis revealed that the number of paraspeckles per nuclei was greater in the WT compared with that in the other mutants, although the expression levels of NEAT1_2 were similar in all the cell lines (Figures 5E and S4C). The calculated numbers of NEAT1_2 per paraspeckle were two to threefold higher in the Δ3′, Δ5′, and Δ5′/Δ3′ mutants than in the WT cells, which was in agreement with the sizes of the paraspeckles as shown in Figure 5C and 5D. Furthermore, EM analysis showed that the number of paraspeckles per cluster in the deletion mutant cells was fewer than that in the WT cells (Figures 5G and S4D). Approximately 79%, 60%, and 87% of the paraspeckles in the Δ3′, Δ5′, and Δ5′/Δ3′ mutant cells, respectively, were present as a single entity. The deletion of the 5′ and/or 3′ terminal domains, corresponding to the A and C blocks, led to enlargement of the paraspeckles through incorporation of more NEAT1_2 transcripts into the paraspeckles, which is consistent with our theoretical predictions.

The block copolymer theory also predicted that the internal organization of NEAT1_2 will influence the shape of the paraspeckles. When the repulsive interactions are reduced or diminished, the condensates form more spherical shapes. The surface repulsive interactions between the A blocks, and those between the C blocks, are reduced by the redistribution of these blocks into the core, so surface tension becomes the dominant contribution that determines the shape of the condensates, leading to the formation of spherical condensates to minimize the surface free energy (Doi, 2013). Our circularity analyses based on the EM observations revealed that the shapes of the paraspeckles in the Δ5′/Δ3′ mutant cells, which have random internal NEAT1_2 organization, were closer to spherical than those in the WT and the other mutants, as theoretically predicted (Figures 5H). These data suggested that the Δ5′/Δ3′ mutant paraspeckles appear to lack most of the A and C blocks, leading to the formation of spherical paraspeckles with random internal organization, similar to macroscopic phase separation.

### The transcriptional level of NEAT1_2 is a key determinant for the size and organization of the paraspeckles

We observed that the paraspeckles in the Δ5′/Δ3′ mutant cells formed large spherical condensates with disordered internal structures. This situation is similar to our theoretical model of arcRNA-driven phase separation using an extension of the Flory-Huggins theory (Yamamoto et al., 2020a). This model predicts that the size of the condensates will become larger with increasing NEAT1_2 expression levels (Figure 6A). To test this prediction experimentally, we examined the sizes of the paraspeckles in the Δ5′/Δ3′/ΔPAS mutant cells with different NEAT1_2 expression levels. We measured the Sx values of the paraspeckles by SRM in changing the NEAT1_2 expression levels using various concentrations of a transcriptional activator of NEAT1_2, the proteasome inhibitor MG132, while MG132 (5 μM) was used in previous experiments as shown in Figures 1–3 (Hirose et al., 2014). The NEAT1_2 levels increased in a MG132 dose-dependent manner (Figure 6B). As the expression levels increased, the paraspeckles in the mutant cell line formed larger spheres with larger Sx values (Figure 6C and 6D). In contrast, as the expression levels increased, the paraspeckles in the WT cells were elongated with almost constant Sx values, as has been reported previously for HeLa cells (Hirose et al., 2014) (Figure 6B–D). These data suggested that, unlike in the WT cells, the size of the spherical paraspeckles in the Δ5′/Δ3′/ΔPAS mutant cells increased by coalescence as long as NEAT1_2 RNPs were available because of the loss of surface repulsive interactions; thus, the size was determined primarily by the NEAT1_2 expression level, which is similar to the situation with macroscopic phase separation.

**Figure 6.**
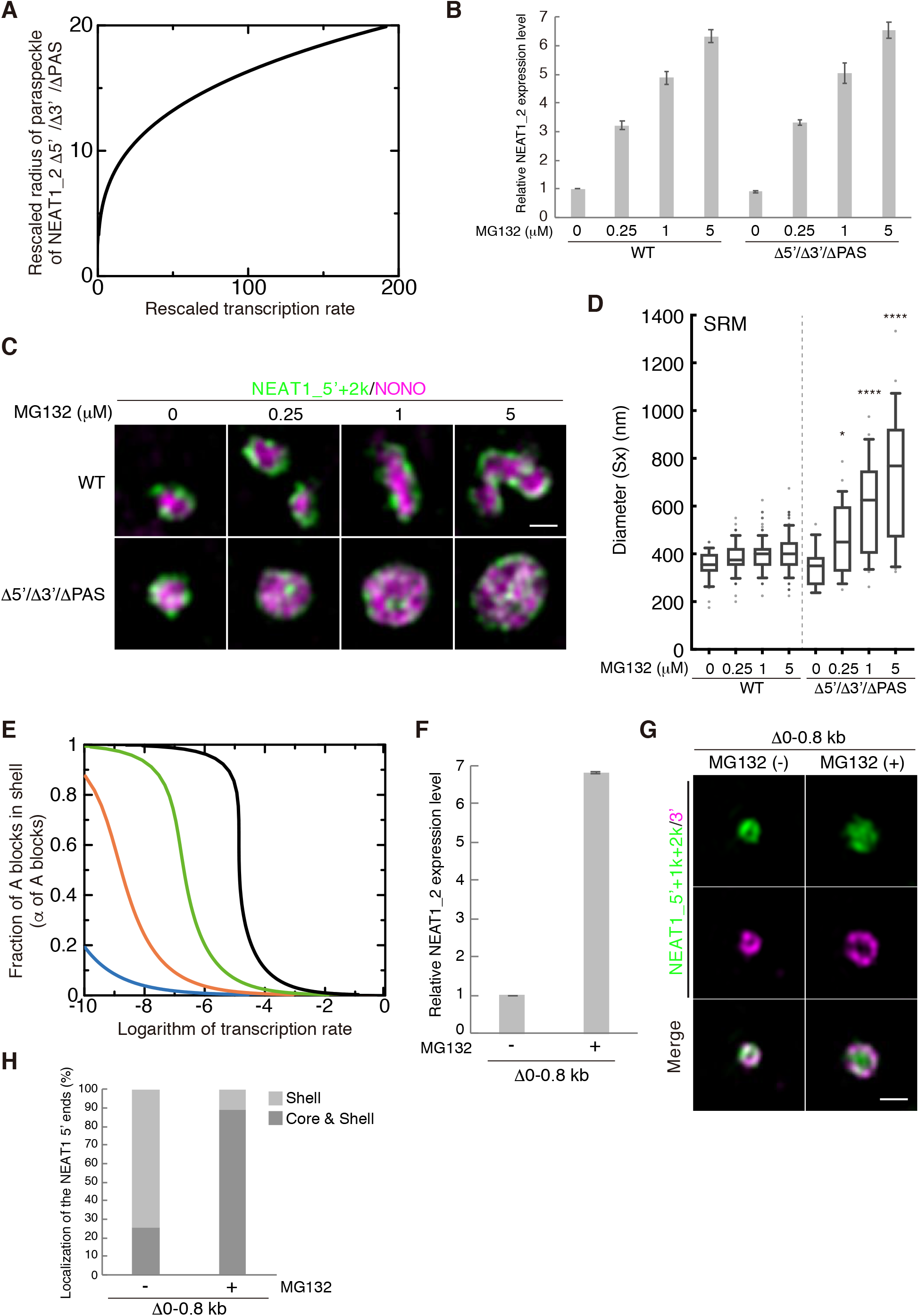
The transcription level of NEAT1_2 is a key determinant for the size and organization of paraspeckles. (A) Theoretical calculation of the rescaled radius of paraspeckles versus the rescaled transcription rate for the case in which the A and C blocks were deleted by CRISPR/Cas9 in the steady state. The graph is derived by assuming that NEAT1_2 is produced at a constant rate. The produced NEAT1_2 RNPs diffuse in the solution and the free diffusion is hindered by the attractive interactions between NEAT1_2 RNPs with the interaction parameter *χ* (we used *χ* = 1.0). NEAT1_2 was degraded at the constant rate *k*_0_. The radius was rescaled by the length scale 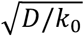 and the transcription rate was rescaled by the inverse time scale *k*0(*D*/*k*0)^3/2^/ν0. (B) Quantitation of NEAT1_2 by RT-qPCR in WT and Δ5′/Δ3′ cells with or without MG132 treatment (6 h). Data are represented as mean ± SD (n = 3). (C) The paraspeckles in WT and Δ5′/Δ3′/ΔPAS cells detected by SRM using NEAT1_5′ and 2k FISH probes (green), and NONO IF (magenta) under the conditions shown in (B). Scale bar, 500 nm. (D) Quantitation of the diameter (Sx) observed by SRM in WT and Δ5′/Δ3′ cells under the conditions shown in (B). WT [mean size: MG132 (–), 354.2 nm (n = 32); MG132 0.25 μM, 380.5 nm (n = 68); MG132 1 μM, 394.1 nm (n = 108); MG132 5 μM, 409.6 nm (n = 103)], Δ5′/Δ3′ [mean size: MG132 (–), 344.4 nm (n = 18); MG132 0.25 μM, 472.1 nm (n = 27); MG132 1 μM, 601.4 nm (n = 27); MG132 5 μM, 721.2 nm (n = 26)]. (*: *P* = 0.029, ****: *P* < 0.0001, compared with WT). (E) Theoretical calculation of the fraction of A (or C) blocks in the shell vs the logarithm of the transcription rate (rescaled by the rate at which the transcripts are spontaneously incorporated in the paraspeckle). The numbers of segments of A blocks were 5.0 (cyan), 7.5 (orange), 10.0 (green), and 12.5228 (black). The parameters used for the calculations are shown in the STAR Methods. (F) Quantitation of NEAT1_2 by RT-qPCR in Δ0–0.8 kb cells with or without MG132 treatment (5 μM for 6 h). Data are represented as mean ± SD (n = 3). (G) The paraspeckles detected by SRM with NEAT1_5′, 1k, and 2k (green), and NEAT1_3′ (magenta) FISH probes in the Δ0–0.8 kb cells under the conditions shown in (H). Scale bar, 500 nm. (H) Graph showing proportion of paraspeckles with localization of the NEAT1 5′ ends to the core and shell or the shell in Δ0–0.8 kb cells under the conditions shown in (E) [MG132 (–): n = 78, MG132 (+): n = 127].

In the block copolymer micelle model, the transcription level of NEAT1_2 is one of the most important parameters controlling the internal organization of NEAT1_2 in the paraspeckles (Yamamoto et al., 2020b). Our theory predicts that as the transcription rate increases, the fractions of the 5′ and 3′ terminal regions that are localized in the shell will decrease (Figure 6E), and these regions are redistributed to the core or become randomly distributed. This redistribution is because increasing the number of NEAT1_2 molecules in the paraspeckle has a greater effect on the repulsive interactions between the A blocks (and those between the C blocks) in the shell compared with the repulsive interactions between the B blocks and A or C blocks in the core (Yamamoto et al., 2020b). To experimentally test this prediction, we observed the paraspeckles in Δ0–0.8 kb mutant cells in the absence and presence of MG132. MG132 treatment strongly enhanced the expression (~7-fold) of NEAT1_2 in the Δ0–0.8 kb mutant cells (Figure 6F). In the absence of MG132, most of the paraspeckles showed shell localization of the 5′ ends as predicted, whereas, in the presence of MG132, most of the paraspeckles showed random localization of the NEAT1 5′ regions (Figure 6G and H), these results were consistent with the data shown in Figure 2B and C. Collectively, our triblock copolymer micelle model of paraspeckles can explain the experimentally observed characteristics of paraspeckles.

## Discussion

We identified the 5′ and 3′ terminal domains of NEAT1_2 arcRNA as the shell-forming domains that determine the spatial organization of NEAT1_2 within paraspeckles (Figure 7A). These shell-forming domains are distinct RNA domains that are located outside of the major assembly domain of NEAT1_2 (8–16.6 kb) (Yamazaki et al., 2018), the deletion of which did not affect the paraspeckle assembly itself. Paraspeckles possess characteristic cylindrical shapes with restricted Sx values and are distinct from other cellular bodies formed by macroscopic phase separation, such as LLPS (A and Weber, 2019; Yamazaki et al., 2019). Thus, paraspeckles are most likely formed through microphase separation that dictates an optimal structure and size of the assemblies (Bates and Bates, 2017; Mai and Eisenberg, 2012; Moughton et al., 2012). Our theoretical approach that treats paraspeckles as amphipathic ABC triblock copolymer micelles could explain and predict the spatial organization of NEAT1_2 within paraspeckles in WT and 5′ and/or 3′ terminal deletion mutants, and how changes in the spatial organizational influenced the size, number, and shape of the paraspeckles (Figure 7B). The model also predicted that the transcription level of NEAT1_2 will influence the size of the paraspeckles and the internal NEAT1_2 organization (Figure 7B). All the predictions were experimentally supported by SRM and EM analyses using the NEAT1_2 deletion mutants. Our experimental and theoretical data suggested that the presence of the 5′ and 3′ domains of NEAT1_2 switch the paraspeckle formation process from macroscopic phase separation to microphase separation. Thus, we propose that the paraspeckles are constructed as block copolymer micelles through microphase separation (Figure 7B).

**Figure 7.**
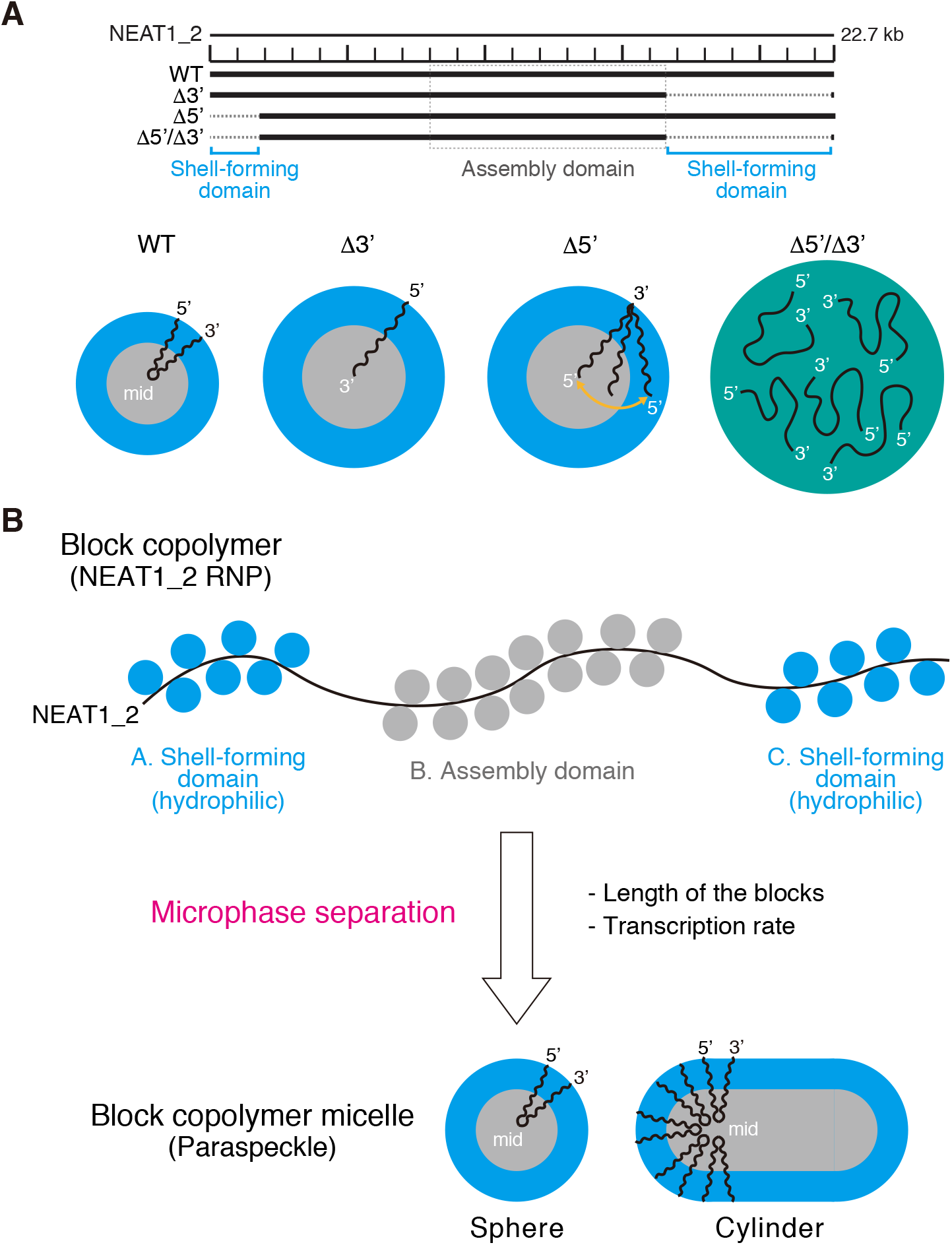
Summary of the block copolymer micelle model of paraspeckles. (A) Functional NEAT1_2 RNA domains for paraspeckle assembly and shell-localization of the 5′ and 3′ ends of the NEAT1_2 within the paraspeckles. Summary of the NEAT1_2 organization within the paraspeckle and the size of the paraspeckles in the representative NEAT1_2 mutants used in this study. (B) The triblock copolymer micelle model for paraspeckles. NEAT1_2 RNPs are treated as ABC triblock copolymers. Putative RNA-binding proteins (blue circles) interact with the shell-forming domains and the proteins involved in the paraspeckle assembly (e.g., NONO) (gray circles) mainly bind to the middle assembly domain. The NEAT1_2 RNPs are microphase-separated into paraspeckles as block copolymer micelles to form spherical and cylindrical shapes. The length of the blocks and the transcription rate of NEAT1_2 influence the organization, size, and shape of the paraspeckles.

In our theoretical model, we treated NEAT1_2 RNPs as triblock copolymers (Figure 7B). lncRNAs usually interact with RBPs to perform their functions and NEAT1_2 has various partner RBPs, such as NONO, which is a major partner protein of the middle assembly domain (the B block) (Nakagawa et al., 2018; Yamazaki et al., 2018). Thus, it is likely that proteins interacting with the 5′ and 3′ shell-forming domains determine the hydrophilic nature of these domains and thus, the localization of these domains in the shell (Figure 7B). In polymer physics, polymer blocks are hydrophilic when the mixing entropy, that is, the contribution of thermal fluctuations, dominates the block–block, water–water, and water–block interactions (Doi 1996). In such cases, the hydrophilic blocks tend to be mixed with the water molecules and the interaction between hydrophilic blocks is considered as effective repulsion (Doi 1996). Further investigations, including an analysis of how the partner proteins of the shell-forming domains generate repulsive interactions, would advance our understanding of the mechanism of paraspeckle structural formation.

In the current theoretical model, for simplicity we assumed that the A and C blocks were composed of identical units, as is the case for uniform polymers, such as synthetic polymers. This would be the situation if the RBPs uniformly interacted with the NEAT1_2 A and C blocks. However, it is plausible that the RBPs bind to specific sub-regions of the NEAT1_2 A and C blocks. Our deletion analyses suggest that the 0–0.8 kb and 16.6–20.2 kb regions of NEAT1_2 are sub-domains that play major roles in the shell localization of the 5′ and 3′ terminal domains, respectively, although we do not rule out the possibility that NEAT1_2 regions outside of these sub-domains also have roles in the shell localization. It has been previously reported that in a mini-NEAT1 mutant that consisted of 0–1 kb and middle assembly (8–16.6 kb) domains and the triple helix structure of NEAT1_2, the 5′ domain was localized in the shell of paraspeckles, suggesting that the 0–1 kb domain was sufficient for shell localization, which is consistent with the results of the present study (Yamazaki et al., 2018). In the Δ16.6–20.2 kb mutant, the 3′ ends were localized in both the core and shell of the paraspeckle, whereas in the Δ3′ mutant, the 3′ ends were localized in the core. One interpretation of these results is that in the Δ16.6–20.2 kb mutant, the 20.2–22.6 kb regions can fluctuate between the core and shell of the paraspeckle. In contrast, the 3′ ends are localized in the core because the Δ3′ mutant lacks the 3′ terminal region that can fluctuate between the core and shell. This interpretation can be applied to the Δ0–0.8 kb and Δ5′ mutants. Thus, in such cases, while the current theoretical model incorporates important aspects of paraspeckle formation, it would be of interest to extend this theoretical model to reflect this fluctuation to model the paraspeckle formation more precisely.

Paraspeckles are often found as clusters (Hirose et al., 2014; Visa et al., 1993) (Figure 5G). This phenomenon can be explained by our block copolymer micelle model. The repulsive interactions between the A blocks, or between the C blocks, in the shell of the paraspeckles may suppress the coalescence of the paraspeckles. This explanation is consistent with the data that the number of paraspeckles in clusters was considerably decreased in the 5′ and/or 3′ deletion mutants (Figure 5G). Repulsive interactions are also crucial in assembling cylindrical paraspeckles. In the micelle model, the structure of the paraspeckles is determined by minimizing the free energy. When the paraspeckles are enlarged upon prolonged transcriptional activation, cylindrical shapes are favorable because i) the free energy owing to the repulsive interactions between A, or C blocks, in the shells of cylindrical paraspeckles is less than that in spherical paraspeckles and ii) the free energy of the middle region of NEAT1_2 in cylindrical paraspeckles is less than that in spherical paraspeckles (Halperin and Alexander, 1989; Semenov et al., 1995; Zhulina et al., 2005). The latter free energy is the elastic energy caused by the polymer blocks, such as the middle region of NEAT1_2, being stretched as springs toward the center of the paraspeckle core because of their connectivity (Doi 1996; Semenov 1985). A cylindrical shape is therefore stable above a certain threshold of the NEAT1_2 expression level, which is consistent with the frequent observations of cylindrical paraspeckles in MG132-treated human, mouse, and opossum cells (Hirose et al., 2014; Yamazaki et al., 2018; Cornelis et al., 2016). In addition to the WT paraspeckles, the paraspeckles in Δ3′ mutant cells often showed elongated shapes (Figures 5H and S4A). The mechanism for the sphere–cylinder morphological transition can be explained by the theory of micelles of AB block copolymers (Halperin and Alexander, 1989; Semenov et al., 1995; Zhulina et al., 2005), although we need to extend our theory to understand the transition from a sphere to a cylinder more quantitatively.

In the present study, we determined that the 5′ and 3′ terminal regions contained shell-forming domains that were independent of the localization of the other region. A previous study has suggested that NEAT1_2 RNPs are looped within a paraspeckle (the looped hypothesis) (Souquere et al., 2010). It has also been proposed that RNA-RNA base pairing mediated long-range interactions between the 5′ and 3′ terminal regions of NEAT1_2 that contributed to the looping (Lin et al., 2018). These long-range interactions might also be mediated by proteins. Although the looping alone cannot explain the core-shell structure of paraspeckles and definitive *in vivo* experimental evidence for the long-range interactions is lacking, further investigations may elucidate a possible role of the looping in the NEAT1_2 arrangement within a paraspeckle.

Block copolymers composed of synthetic monomers have been well studied to generate a variety of materials with distinct physical properties and functions, and various nano-order shapes of the self-assemblies, including spheres, cylinders, lamellae, and vesicles, can be formed by changing the length of the blocks (Mai and Eisenberg, 2012; Moughton et al., 2012). In this study, we generated paraspeckles with different sizes and shapes by deleting the NEAT1_2 A and/or C blocks. Thus, it is tempting to speculate that RNAs have the potential to form various structures with distinct properties as polymer scaffolds, likely with RBPs, in cells. Nuclear stress bodies constructed by HSATIII arcRNAs possess a sea-island structure, a typical structure formed by block copolymers, and thus, these bodies might be formed by a similar mechanism (Chiodi et al., 2000; Kawaguchi et al., 2015; Ninomiya et al., 2019). Furthermore, as NEAT1_2 has modular domains, such as UG repeats for TDP-43 recruitment (Modic et al., 2019), it might be possible that addition of such modular domains to arcRNAs may result in the acquisition of new functions by the recruitment of specific RBPs. A similar concept has been shown in block copolymer research (Lutz et al., 2013; Ouchi et al., 2011). Therefore, the domains of arcRNAs may be used to determine the size, structure, and function of condensates, and thus, it may be possible to design RNA-based microphase-separated bodies.

In addition to our theoretical framework for RNA-driven macroscopic phase separation (Yamamoto et al., 2020a), our block copolymer micelle model will be useful for further experimental investigation and theoretical modeling of RNA-driven phase separation. As various molecular functions of paraspeckles have been reported (Bonetti et al., 2020; Cai et al., 2020; Chen and Carmichael, 2009; Hirose et al., 2014; Imamura et al., 2014; Jiang et al., 2017; Li et al., 2017; West et al., 2014; West et al., 2016), using this model may contribute to understanding the relationship between the core-shell structures, the number and size of paraspeckles, and the paraspeckle functions. It has been recently proposed that chromatins, which are long polymers, behave as block copolymers (Belanghzal et al., 2020; Hildebrand and Dekker, 2020) and our work develops the concept that RNPs can function as block copolymers to form highly ordered microphase-separated condensates in cells. Further work is needed to elucidate the molecular details (e.g., the components, including RBPs, and the RNA sequence and structural elements) of the RNP block copolymers. Finally, this work provides a foundation for future studies of microphase-separated RNP block copolymer micelles in cells.

## Acknowledgments

The authors thank C. Fujikawa and A. Kubota for technical support and the members of the Hirose laboratory for valuable discussions. This research was supported by KAKENHI grants from the Ministry of Education, Culture, Sports, Science, and Technology (MEXT) of Japan [to T. Yamazaki (17K15058, 19K06479, 19H05250), T. Yamamoto (19H05259, 18K03558), and TH (26113002, 17H03630, 17K19335, 19K22374, 20H00448, 20H05377)], JST, PRESTO Grant Number JPMJPR18KA (to T. Yamamoto), the Mochida Memorial Foundation for Medical and Pharmaceutical Research (to T. Yamazaki), the Naito Foundation (to T. Yamazaki), the Takeda Science Foundation (to T. Yamazaki), and Tokyo Biochemical Research Foundation (to TH).

## Author contributions

T. Yamazaki, T. Yamamoto, and TH conceived and designed this study. T. Yamazaki and HY conducted the experiments, except for the EM and theoretical analyses. SS and GP performed the EM analyses. T. Yamamoto performed the theoretical analyses. SN technically supported the SRM. T. Yamazaki, T. Yamamoto, GP, and TH wrote the manuscript.

## Declaration of interests

The authors declare no competing interests.

## STAR METHODS

### KEY RESOURCES TABLE

#### CONTACT FOR REAGENT AND RESOURCE SHARING

Further information and requests for resources and reagents should be directed to, and will be fulfilled by, the Lead Contact, Tetsuro Hirose.

#### EXPERIMENTAL MODEL AND SUBJECT DETAILS

##### HAP1 cell line

HAP1 cells (Horizon Discovery) were maintained in IMDM (Gibco) supplemented with 10% FBS purchased from Gibco or Sigma-Aldrich.

#### METHOD DETAILS

##### Genome editing using CRISPR/Cas9

CRISPR-mediated deletions of the *NEAT1* gene were performed as previously described (Yamazaki et al., 2018). The sequences of the sgRNA and HAP1 mutant cell lines established in this study are listed in Supplemental Table S1. The sgRNAs were cloned into a PX330-B/B vector (Yamazaki et al., 2018) for use for the deletions of the *NEAT1* gene.

##### Reverse transcription-quantitative PCR (RT-qPCR)

Purification, reverse-transcription of total RNA, and qPCR for NEAT1_1 and NEAT1_2 were performed as previously described in detail (Yamazaki et al., 2018). The primers used in this study are listed below. For NEAT1_2 detection, the NEAT1-2’ primer set (forward primer: 5′-CAATTACTGTCGTTGGGATTTAGAGTG-3′, reverse primer: 5′- TTCTTACCATACAGAGCAACATACCAG-3′) was used. For detection of NEAT1_2, the NEAT1-6 primer set (forward primer: 5′-CAGTTAGTTTATCAGTTCTCCCATCCA-3′, reverse primer: 5′-GTTGTTGTCGTCACCTTTCAACTCT-3′) or NEAT1-12 primer set (forward primer: 5′-TGTGTGTGTAAAAGAGAGAAGTTGTGG-3′, reverse primer: 5′- AGAGGCTCAGAGAGGACTGTAACCTG-3′) was used. 18S (forward primer: 5′- TTTAAGTTTCAGCTTTGCAACCATACT-3′, reverse primer: 5′- ATTAACAAGAACGAAAGTCGGAGGT-3′) or GAPDH (forward primer: 5′- ATGAGAAGTATGACAACAGCCTCAAGAT-3′ reverse primer: 5′- ATGAGTCCTTCCACGATACCAAAGTT-3′) primer sets were used as loading controls.

##### RNA-FISH and immunofluorescence

RNA-FISH and immunofluorescence were performed as previously described (Yamazaki et al., 2018). Confocal and super-resolution microscopic analyses (structured illumination microscopy: SIM) were performed as previously described (Yamazaki et al., 2018). LSM900 with Airyscan2 for super-resolution imaging was used in the analysis of the quantification of paraspeckle numbers per nucleus. The antibodies used in this study are listed in the Key Resource Table. The NEAT1 FISH probes for SRM analyses against 5′ (NEAT1: +1 to +1000), 1k (+1208 to +1935), 2k (+2025 to +2783), 15k (+15401 to +16612), and 19k (+19401 to +20040) and 3′ (+21743 to +22580) were transcribed as antisense RNAs. smFISH was performed using the human NEAT1_m probe (LGC Biosearch Technologies) as previously described (Yamazaki et al., 2018).

##### Electron microscopy

Ultrastructural studies were carried out on ultra-thin sections of Lowicryl K4M-embedded cell pellets as previously described (Souquere and Pierron, 2015). Duplicated samples of WT and mutant HAP1 cells grown in the absence or presence of 5 μM MG132 for 17 h were fixed *in situ* with 1.6% glutaraldehyde (Electron Microscopic Sciences) or 4% formaldehyde (Electron Microscopic Sciences), scraped off from the plastic containers, and centrifuged. The cell pellets were equilibrated in 30% methanol and deposited in a Leica EM AFS2/FSP automatic reagent handling apparatus (Leica Microsystems). Lowicryl polymerization under UV was performed for 40 h at –20°C and 40 h at 20°C. The DNA probes for high resolution *in situ* hybridization (EM-ISH) were PCR-amplified DNA fragments, biotinylated by nick translation (Roche) with biotin-16-dUTP (Roche) but with no TTP in the reaction mix. The NEAT1_5′ DNA probe (+230–1721 and +1751– 3244), the D2 probe (+12841–14160 and +14735–15897) and hybridization conditions, and detection of RNA/DNA hybrids with goat anti-biotin antibody conjugated to 10 nm gold particles (BBI International) were as previously described (Souquere et al., 2010; Souquere and Pierron, 2015). Occasionally, ultra-thin sections of formaldehyde-fixed cells were pre-treated with protease (0.2 mg/ml) for 15 min at 37°C to enhance access of the biotinylated DNA probes to the NEAT1 targets. The thin sections were briefly contrasted with uranyl acetate and analyzed with a Tecnai Spirit (FEI). Digital images were taken with an SIS Megaview III charge-coupled device camera (Olympus). The geometry of the nuclear body sections [short axes (Sx), long axes (Lx), and surface areas] and gold particle distribution were determined with AnalySIS (Olympus Soft Imaging Solutions).

##### Triblock copolymer micelle model

We have recently developed a model that treats paraspeckles as micelles of triblock copolymers (Yamamoto et al., 2020b, in which the details of the mathematical formalism are described). The construction of the model was motivated by the fact that paraspeckles show a core-shell structure, analogous to polymer micelles. In our theory, NEAT1_2 is treated as an ABC triblock copolymer. The B blocks are localized in the core and the C blocks are localized in the shell. A fraction, *α*, of the A blocks are in the shell and the other fraction, 1 − *α*, are in the core.

Polymer micelle theory predicts that the optimal size of polymer micelles composed of AB block copolymers (B blocks form the core and A blocks form the shell) is determined by the free energy composed of 1) the surface free energy (which is the total surface area of the core multiplied by the surface tension) and 2) the free energy because of the repulsive excluded-volume interactions between the segments of A blocks in the shell (Halperin and Alexander, 1989; Semenov et al., 1995; Zhulina et al., 2005). Many theories regarding polymer micelles assume that all the A blocks are in the shell. In paraspeckles, the A and C blocks in the shell are separated in different domains and a fraction, 1 − *α*, of the A blocks are in the core. Therefore, we took into account 3) the free energy from the repulsive excluded-volume interactions between the C blocks in the shell; 4) the free energy from the repulsive interactions between the B blocks and the A blocks in the core; and 5) the free energy from the mixing entropy (that contributes to equally distributing the A blocks to the shell and the core) in an extension of the free energy calculations for polymer micelles. The free energy because of the conformational entropy of B blocks in the core is small and we neglected this free energy (Halperin and Alexander, 1989; Semenov et al., 1995; Zhulina et al., 2005). We determined the fraction, *α*, of A blocks in the shell by minimizing the free energy, *F*_n_(*α*). With an optimal value of the fraction, *α*, the free energy is a function only of the number *α* of NEAT1_2 molecules.

In contrast to polymer micelles spontaneously assembled in a solution at equilibrium, the assembly of paraspeckles had a strong correlation with the transcription of NEAT1_2. This correlation is probably because RNA-binding proteins already bind to nascent NEAT1_2 transcripts, the production of which is not yet completed, and these nascent transcripts are incorporated one by one into the paraspeckle at a constant rate *k*_tx_. Nascent transcripts are connected to the transcription site via RNA polymerase II and thus, can be incorporated in the paraspeckle at the transcription site without translational entropy cost. The growth of the paraspeckle was analyzed using the Master equation. The Master equation predicts that the number of transcripts in the paraspeckle in the steady state is determined by the minimum of the effective free energy

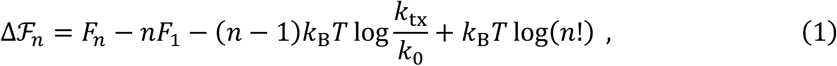

where *k*_B_ is the Boltzmann constant, *T* is the absolute temperature, and *k*_0_ is the rate at which NEAT1_2 transcripts in the solution are spontaneously incorporated in the paraspeckle (by the thermal process). Our model can predict the number *n* of transcripts in a paraspeckle, the fraction *α* of A blocks in the shell, the radius of the paraspeckle in the steady state as a function of the transcription rate *k*_tx_, and the number of segments in the A blocks.

The following parameters were used for the numerical calculations: the number of segments in the B blocks was 40, the number of segments in the C blocks was 20, the dimensionless surface tension was 0.5, the interaction parameter between the A segments and B segments in the core was 1.0, the excluded volume of the A and C blocks was the cubic of the segment length (athermal solvent conditions). In Figure 4D and Figure 5A and B, we used log(*k*_tx_/*k*_0_) = −8.0.

##### Phase separation model

Both the A and C blocks of Δ3′/Δ5′ NEAT1_2 transcripts are either absent or short. The excluded-volume interactions between the A blocks and those between the C blocks are small and thus, the dynamics of paraspeckles composed of such transcripts is governed by the surface tension. Therefore, we used an extension of the Flory-Huggins theory [which is the standard theory of phase separation in a polymer solution (see standard textbooks of polymer physics, such as Doi 1996)] to predict the growth of paraspeckles composed of Δ3′/Δ5′ NEAT1_2 (Yamamoto et al., 2020a, in which the details of the mathematical formalism are described).

In our theory, Δ3′/Δ5′ NEAT1_2 transcripts are produced at the transcription site at a constant rate. These transcripts diffuse in the solution and the diffusion is slowed down by the attractive interactions between transcripts. The magnitude of the attractive interactions is represented by the interaction parameter *χ*. The transcripts degrade at a constant rate. Our theory predicts that when the interaction parameter *χ* is larger than the critical value *χ*_c_ at equilibrium, the local volume fraction of transcripts jumps from a relatively large value to a smaller value at a distance from the transcription site. This distance defines the radius of the paraspeckle. The jump in volume fraction results from the instability because of the phase separation. This theory can predict the radius of the paraspeckle in the steady state as a function of the transcription rate, the diffusion constant, and the degradation rate.

#### QUANTIFICATION AND STATISTICAL ANALYSIS

Volocity (PerkinElmer) was used for quantification of the paraspeckle size and sum intensity of smFISH signals with intensity and size threshold. Fiji software (NIH) was used for quantification of the Sx values (distance between the peak and peak) of the paraspeckles observed by SRM using a plot profile (West et al., 2016). The sizes of paraspeckles observed by EM were determined with AnalySIS (Olympus Soft Imaging Solutions). Prism 7 software (GraphPad) was used for graphing and statistical analysis. Each box plot shows the median (inside line), 25–75 percentiles (box bottom to top), and 10–90 percentiles (whisker bottom to top). Kruskal-Wallis test with Dunn’s multiple comparison test was used for Figures 5C, 5D, 5E, 5H, 6D, S1C, S2B, S3B, S4A, and S4B. Mann-Whitney test (two-tailed) was used for Figure S2D.

#### DATA AND CODE AVAILABILITY

The scripts used for the theoretical modeling can be found at https://figshare.com/s/be1d03f274642ed1416a.

## Supplemental Figures

**Figure S1.**
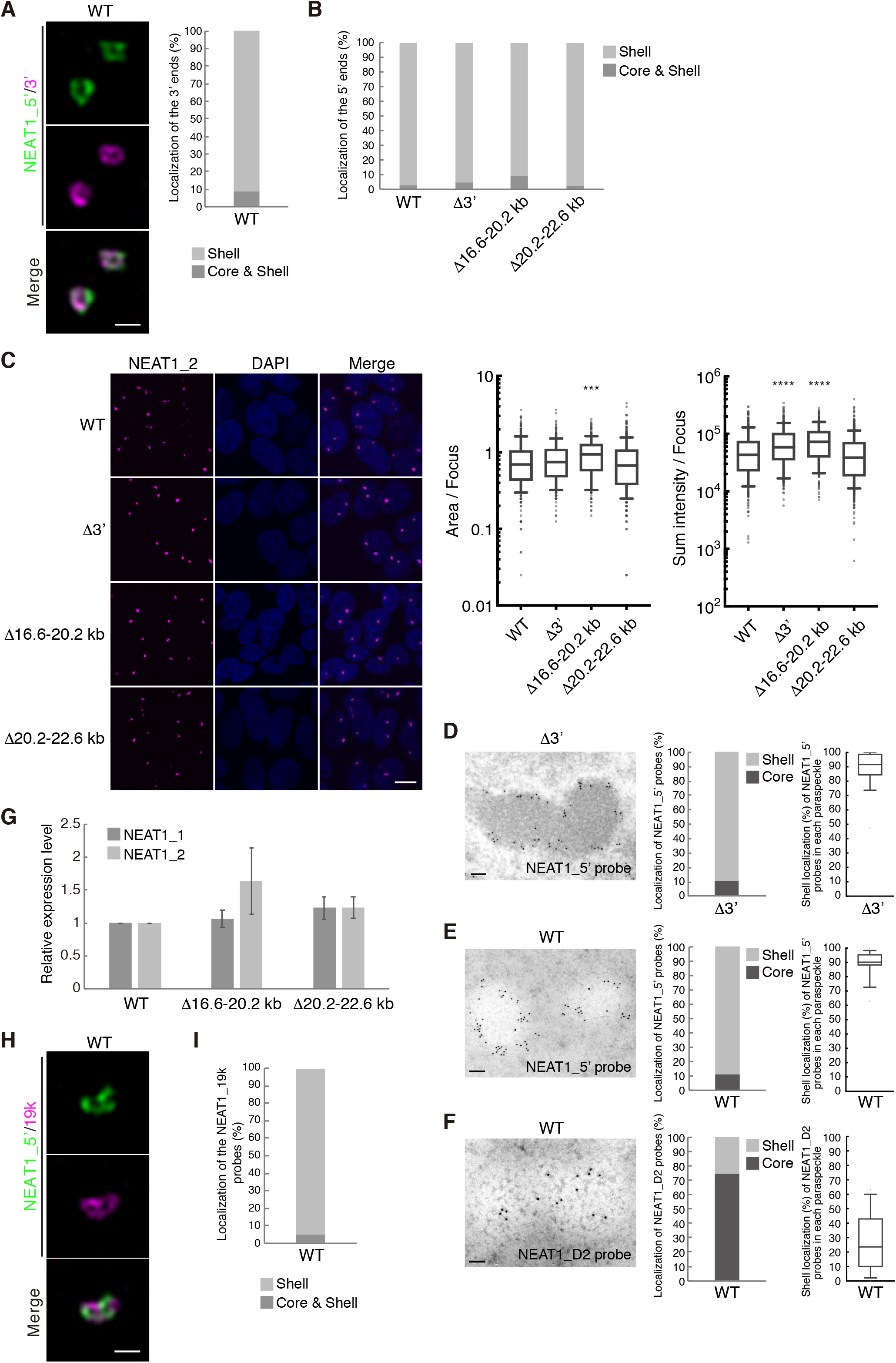
Characterization of NEAT1 mutants with deletions in the NEAT1_2 3′ terminal regions, related to Figure 1. (A) (left) SRM images of the paraspeckles in HAP1 WT cells detected by NEAT1_5′ (green) and NEAT1_3′ (magenta) FISH probes with MG132 treatment (5 μM for 6 h). Scale bar, 500 nm. (right) Graph showing the proportion of paraspeckles with localization of the NEAT1 3′ ends to the core and shell or the shell in WT cells (n = 167). (B) Graph showing the proportion of paraspeckles with localization of the NEAT1 5′ ends to the core and shell or the shell in WT, Δ3′, Δ16.6–20.2 kb, and Δ20.2–22.6 kb cells treated with MG132 (5 μM for 6 h) (WT: n = 115, Δ3′: n = 21, Δ16.6-20.2 kb: n = 161, and Δ20.2-22.6 kb: n = 89). (C) (left) Detection of NEAT1_2 by single-molecule FISH (smFISH) (magenta) in HAP1 WT, Δ3′, Δ16.6–20.2 kb, and Δ20.2–22.6 kb cells treated with MG132 (5 μM for 6 h). Nuclei were stained with DAPI. Scale bar, 10 nm. (right) Quantitation of area and sum intensity per paraspeckle in each cell line. (***: *P* = 0.0001, ****: *P* < 0.0001, compared with WT) (D) (left) EM observations of the paraspeckles in MG132-treated (5 μM for 17 h) HAP1 Δ3′ cells using NEAT1_5′ probe. Scale bar, 100 nm. (middle) Graph showing the proportion of the localization of NEAT1_5′ probes (838 gold particles) within the paraspeckles in Δ3′ cells. (right) Graph showing the proportion of localization of NEAT1_5′ probes in each paraspeckle in Δ3′ cells (n = 21). (E) (left) EM observation of the paraspeckles in MG132-treated (5 μM for 17 h) HAP1 WT cells using NEAT1_5′ probe. Scale bar, 100 nm. (middle) Graph showing the proportion of localization of NEAT1_5′ probes (601 gold particles) within the paraspeckles in WT cells. (right) Graph showing the proportion of localization of NEAT1_5′ probes in each paraspeckle in WT cells (n = 16). (F) (left) EM observation of the paraspeckles in MG132-treated (5 μM for 17 h) HAP1 WT cells using the NEAT1_D2 probe. Scale bar, 100 nm. (middle) Graph showing the proportion of localization of NEAT1_D2 probes (461 gold particles) within the paraspeckles in WT cells. (right) Graph showing the proportion of localization of NEAT1_D2 probes in each paraspeckle in WT cells (n = 24). (G) Quantitation of the relative expression levels of NEAT1_1 and NEAT1_2 by RT-qPCR in HAP1 WT, Δ3′, Δ16.6–20.2 kb, and Δ20.2–22.6 kb cells treated with MG132 (5 μM for 6 h). Data are represented as mean ± SD (n = 3). (H) SRM images of the paraspeckles in MG132-treated (5 μM for 6 h) WT cells detected by NEAT1_5′ (green) and NEAT1_19k (magenta) FISH probes. Scale bar, 500 nm. (I) Graph showing the proportion of paraspeckles with localization of the NEAT1_19k probes to the core and shell or the core in WT cells treated with MG132 (5 μM for 6 h) (n = 103).

**Figure S2.**
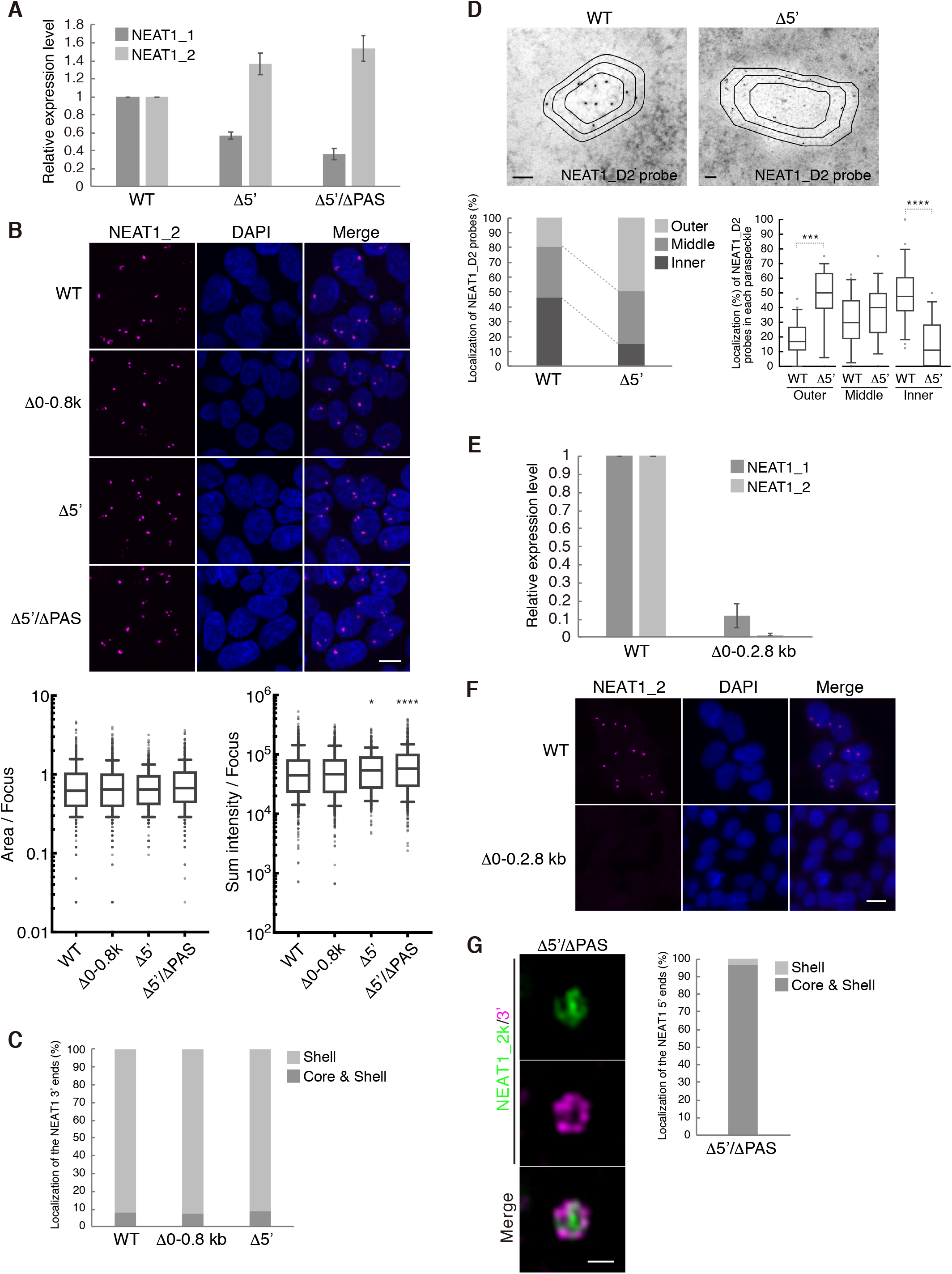
Characterization of NEAT1 mutants with deletions in the NEAT1_2 5′ terminal regions, related to Figure 2. (A) Quantitation of the relative expression levels of NEAT1_1 and NEAT1_2 by RT-qPCR in HAP1 WT, Δ5′, and Δ5′/ΔPAS cells treated with MG132 (5 μM for 6 h). Data are represented as mean ± SD (n = 3). (B) (upper) Detection of NEAT1_2 by smFISH (magenta) in WT, Δ0–0.8 kb, Δ5′, and Δ5′/ΔPAS kb cells treated with MG132 (5 μM for 6 h). Nuclei were stained with DAPI. Scale bar, 10 nm. (lower) Quantitation of area and sum intensity per paraspeckle in each cell line. (*: *P* = 0.0420, ****: *P* < 0.0001, compared with WT). (C) Graph showing the proportion of paraspeckles with localization of the NEAT1 3′ ends to the core and shell or the shell in WT, Δ0–0.8 kb, and Δ5′ cells treated with MG132 (5 μM for 6 h) by SRM analyses (WT: n = 167, Δ0–0.8 kb: n = 77, Δ5′: n = 48). (D) (upper panels) EM observations of the paraspeckles in MG132-treated (5 μM for 17 h) WT and Δ5′ cells using the NEAT1_D2 probe. (bottom, left) Graph showing the proportion of localization (three layers: outer, middle, and inner layers) of NEAT1_D2 probes (WT: 307 gold particles, Δ5′: 167 gold particles) within the paraspeckles in HAP1 Δ5′ cells. Scale bar, 100 nm. (bottom, right) Graph showing the proportion of localization of NEAT1_D2 probes in each paraspeckle in WT (n = 22) and Δ5′ cells (n = 15). (***: *P* = 0.0001, ****: *P* < 0.0001, compared with WT). (E) Quantitation of the relative expression levels of NEAT1_1 and NEAT1_2 by RT-qPCR in HAP1 WT and Δ0–2.8 kb cells. Data are represented as mean ± SD (n = 3). (F) Detection of the paraspeckles by smFISH in WT and Δ0–2.8 kb cells. Scale bar, 10 nm. (G) (left) The paraspeckles in Δ5′/ΔPAS cells treated with MG132 treatment (5 μM for 6 h) detected with SRM by NEAT1_2k (green) and 3′ (magenta) FISH probes. Scale bar, 500 nm. (right) Graph showing the proportion of paraspeckles with localization of the NEAT1 5′ ends to the core and shell or the shell in Δ5′/ΔPAS cells (n = 31).

**Figure S3.**
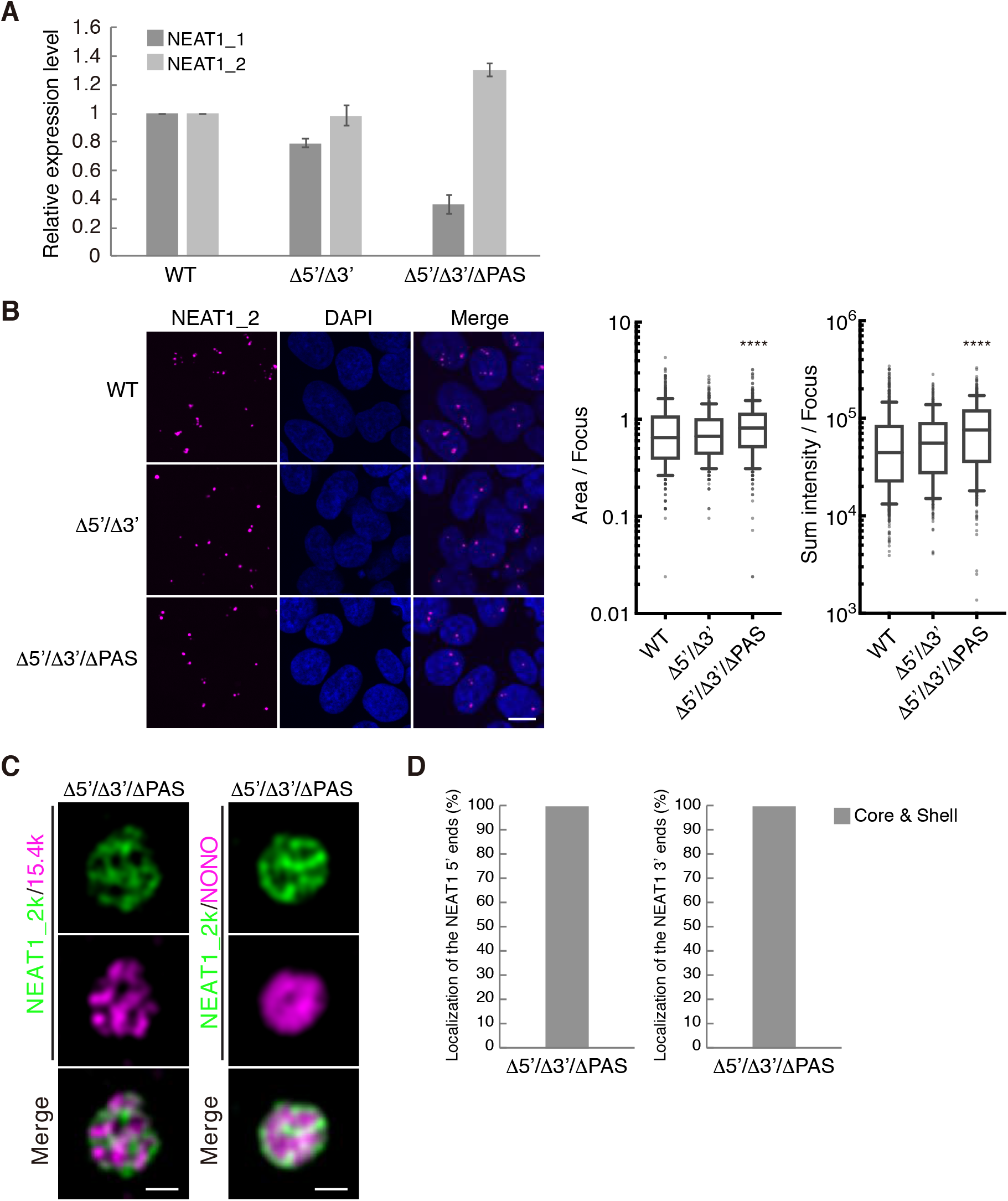
Characterization of NEAT1 mutants with deletions in both the NEAT1_2 5′ and 3′ regions, related to Figure 3. (A) Quantitation of the relative expression levels of NEAT1_1 and NEAT1_2 by RT-qPCR in HAP1 WT, Δ5′/Δ3′, and Δ5′/Δ3′/ΔPAS cells treated with MG132 (5 μM for 6 h). Data are represented as mean ± SD (n = 3). (B) (left) Detection of NEAT1_2 by smFISH (magenta) in WT, Δ5′/Δ3′, and Δ5′/Δ3′/ΔPAS cells treated with MG132 (5 μM for 6 h). Nuclei were stained with DAPI. Scale bar, 10 nm. (right) Quantitation of area and sum intensity per paraspeckle in each cell line. (****: *P* < 0.0001, compared with WT). (C) The paraspeckles detected with SRM by NEAT1_2k FISH probe (green), and NEAT1_15k FISH probe or NONO IF (magenta) in the Δ5′/Δ3′/ΔPAS cells treated with MG132 (5 μM for 6 h). Scale bar, 500 nm. (D) Graph showing proportion of paraspeckles with localization of the NEAT1 5′ (left, n = 41) and 3′ (right, n = 14) ends to the core and shell or the shell in Δ5′/Δ3′/ΔPAS cells treated with MG132 (5 μM for 6 h).

**Figure S4.**
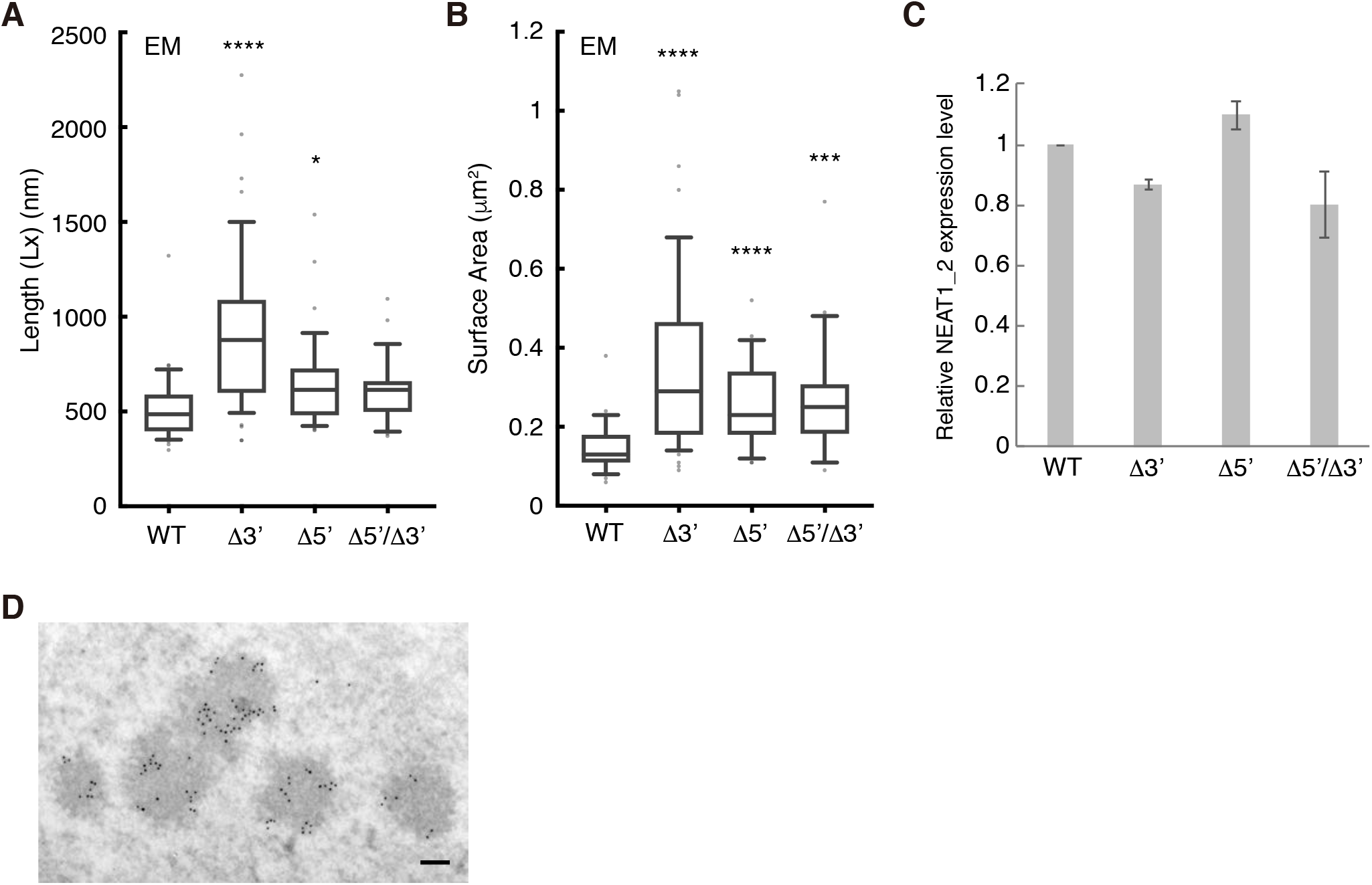
Additional size and structure parameters determined by EM for the paraspeckles of NEAT1 mutants, related to Figure 5. (A) Length (Lx) of the paraspeckles determined by EM in HAP1 WT, Δ3′, Δ5′, and Δ5′/Δ3′ cells treated with MG132 (5 μM for 17 h). WT (mean size: 518.6 nm, n = 35), Δ3′ (mean size: 937.1 nm, n = 49), Δ5′ (mean size: 648.1 nm, n = 39), and Δ5′/Δ3′ (mean size: 608.9 nm, n = 28). (*: *P* = 0.0451, ****: *P* < 0.0001, compared with WT). (B) Surface area of the paraspeckles determined by EM in WT, Δ3′, Δ5′, and Δ5′/Δ3′ cells treated with MG132 (5 μM for 17 h). WT (mean area: 0.1497 μm^2^, n = 35), Δ3′ (mean area: 0.3684 μm^2^, n = 49), Δ5′ (mean area: 0.2582 μm^2^, n = 39), and Δ5′/Δ3′ (mean area: 0.2657 μm^2^, n = 28). (***: *P* = 0.0008, ****: *P* < 0.0001, compared with WT). (C) An analysis of the NEAT1_2 expression levels for quantification of the NEAT1_2 number per paraspeckle. (D) A representative image of a cluster of paraspeckles detected by paraspeckle marker proteins, BRG1 (gold particles) under EM in the presence of MG132. In this image, five paraspeckles form a cluster. Scale bar, 100 nm.

---

**Table S1.**
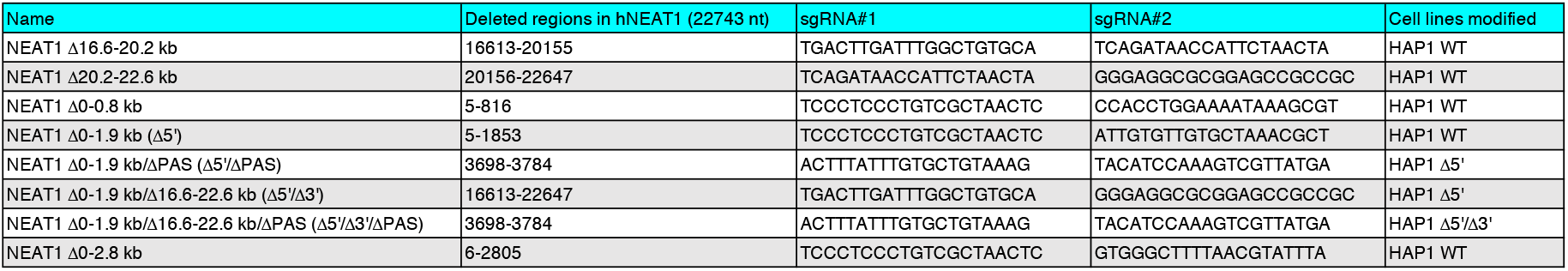
HAP1 mutant cell lines established in this study, Related to STAR Methods

## Notes

### Competing Interest Statement

The authors have declared no competing interest.

